# Multi-millennial genetic resilience of Baltic diatom populations disturbed in the past centuries

**DOI:** 10.1101/2025.03.10.642313

**Authors:** Alexandra Schmidt, Sarah Bolius, Anna Chagas, Juliane Romahn, Jérôme Kaiser, Helge W. Arz, Miklós Bálint, Anke Kremp, Laura S. Epp

**Author notes:** Correspondence Alexandra Schmidt, University of Konstanz, Mainaustr. 252, 78464 Constance, Germany, Laura S. Epp, University of Konstanz, Mainaust. 252, 78464 Constance, Germany.

## Abstract

Little is known about the genetic diversity and stability of natural populations over millennial time scales, although the current biodiversity crisis calls for heightened understanding. Marine phytoplankton, the primary producers forming the basis of food webs in the oceans, play a pivotal role in maintaining marine ecosystems health and serve as indicators of environmental change. This study examines the genetic diversity and shifts in allelic composition in the diatom species *Skeletonema marinoi* over ∼ 8000 years in the Baltic Sea by analyzing chloroplast and mitochondrial genomes. Ancient environmental DNA (aeDNA) from sediment cores demonstrates stability and resilience of genetic composition and diversity of this species across millennia in the context of major climate events. Accelerated change in allelic composition is observed from historical periods onwards, coinciding with times of intensifying human activity, like the Roman Empire, the Viking Age, and the Hanseatic Age, suggesting that anthropogenic stressors have profoundly impacted this species for the last two millennia. The data indicate a very high natural stability and resilience of the genomic composition of the species and underscore the importance of uncovering genomic disruptions caused by human impact on organisms, even those not directly exploited, to better predict and manage future biodiversity.

## Introduction

Human activities have had a profound impact on natural environments, resulting in the endangerment of a multitude of ecosystems, including marine environments^1^. Phytoplankton forms the basis of marine food webs, and, as crucial components of marine ecosystems, changes in this group are central to understanding ecosystem shifts^2^. Among the phytoplankton, diatoms, which play a significant role in global biogeochemical cycles, are sensitive bioindicators^3^, as the different species have specific requirements. *Skeletonema marinoi*, a prominent marine diatom species^4^, is influenced by temperature, migration, and human activities^5^. For this species, changes in bloom timing^6^ and optimal growth temperature^7^ have already been observed as a result of recent climate change. Investigation of marine phytoplankton response to current global changes are mostly based on taxonomic diversity, but recent studies show that diatoms also quickly adapt to environmental changes through acclimation and genetic adaptation on a population genomic level^8^. In fact, intraspecific genomic variation is an important factor in the response and resilience of diatoms to environmental perturbations^9^. Thus, investigating the genomic composition of diatom populations and its changes is crucial for understanding their adaptability to environmental changes.

The Baltic Sea, despite its relatively brief history, has undergone significant transformations, making it suitable for studies addressing organism responses to environmental changes. For the last ∼ 10,000 years, these transformations include a transition between fresh and brackish water around 8,500-8,000 years ago^10^ and major climate changes since then^11^. Due to its shallow enclosed nature and a steep salinity gradient, modern biodiversity is relatively low^12^. Current pressures such as pollution, eutrophication and global warming present significant challenges to the biota^13^.

Sediments can serve as ecological archives when undisturbed, offering insights into past environmental changes and biodiversity shifts^14^, and provide long time series of phytoplankton dynamics. Phytoplankton archives in sediments include biomarkers, microfossils, (living) resting stages, and sedimentary ancient DNA (sedaDNA). These remains contain information on biodiversity and can reveal changes in adaptive traits^7,15^. Here, we develop a time series of population-level responses of the diatom *S. marinoi* in the Baltic Sea, by employing a targeted approach to enrich chloroplast and mitochondrial genomes using hybridization enrichment. This allows us to analyze genetic diversity and shifts in allelic composition across complete organellar genomes. We investigate 1) the degree of stability, response and resilience of *S. marinoi* to natural environmental changes in the Baltic Sea over the last ca. 8000 years, and 2) identify changes in the genetic composition of populations in recent centuries of heightened anthropogenic activities.

## Results

We analyzed sediment samples of two cores located in the Baltic Sea, the Eastern Gotland Basin (EGB) and the Gulf of Finland (GOF) (Fig. 1A), covering a time span up to approximately the Ancylus Lake (extrapolated age: 8148 cal yr BP) (BP; with present = 1950 Common Era) (Fig. 1B). We investigated the population structure of *S. marinoi* over this period at the two locations by focusing on Single Nucleotide Polymorphisms (SNPs) on the organelle genomes (Fig. 1C). This was achieved by target enrichment of the chloroplast and the mitochondrion.

**Fig. 1.**
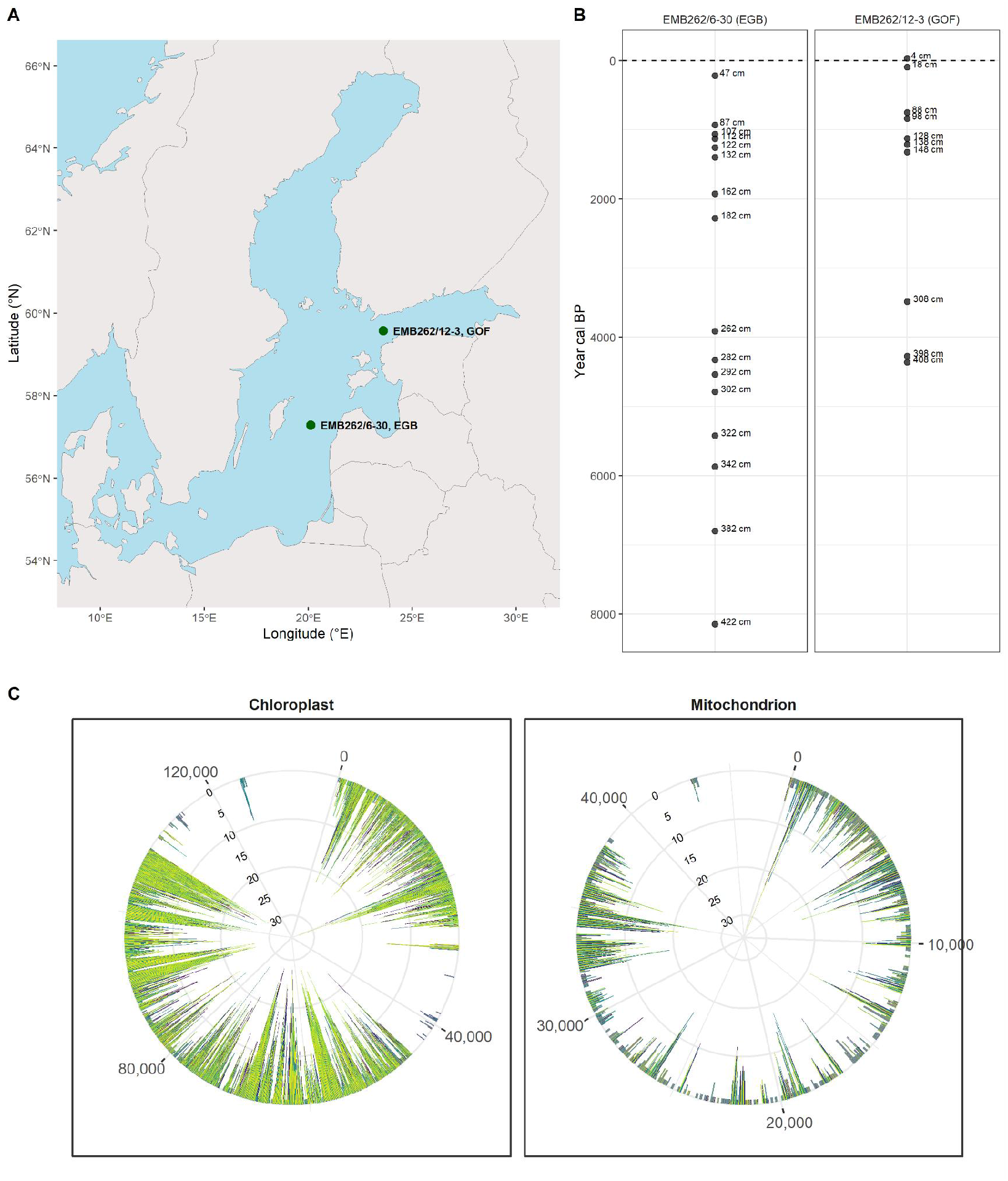
A) Location of the coring sites and corresponding cores in the central Baltic Sea. Gulf of Finland (GOF; core EMB262/12-3GC) and Eastern Gotland Basin (EGB; core EMB262/6-30GC). B) Sample distribution (in cm) in the sediment cores and corresponding ages. C) Number of SNPs called per 100 base pairs across the genome for each sample. The mitochondrion and chloroplast genomes of *S. marinoi* are shown. The gaps indicate the repeat masked regions.

This study provides an analysis of the genetic dynamics of *S. marinoi* populations in the EGB and GOF over millennia, revealing insights into the effects of both natural environmental changes and anthropogenic activities. The use of RNA baits for hybridization enrichment resulted in a higher degree of resolution, enabling the identification of a greater number of SNPs in both mitochondrial and chloroplast genomes. The analysis revealed that climate events exert an influence on genetic variations, with specific climate events aligning with temporal variations in the population’s genetic makeup, but that the genetic composition reverted back to a state that was mostly stable across millennia in between these events. Longer lasting patterns of genetic change over time were observed at both sites in the past two millennia, coinciding with periods of intensified anthropogenic activities.

These findings imply high resilience of *S. marinoi* populations across time and climate events across millennia, along with more recent changes.

## General data assessment

### DNA vs. RNA baits

In order to study past *S. marinoi* population dynamics in the EGB and GOF, we employed DNA and RNA baits for hybridization enrichment of full chloroplast and mitochondrial genomes. The use of RNA baits resulted in a higher degree of resolution. Following trimming, DNA and RNA baits yielded 34.9 million and 51.8 million sequences, respectively. Of these, 34.8 million (DNA) and 35.3 million (RNA) reads were successfully mapped to reference organelles (Fig. 2A and Supplementary Table S1). In the case of RNA baits, 6.41 million reads were mapped to the mitochondrion, while 28.9 million were mapped to the chloroplast. For DNA baits, the numbers were 6.27 million and 28.6 million, respectively. The RNA baits identified a greater number of SNPs in both the mitochondrial (301 vs. 92) and chloroplast (1716 vs. 403) genomes. Consequently, the analysis focuses on the results obtained from RNA baits. The results of the DNA bait analysis can be found in the supplementary section 2.1.

**Fig. 2:**
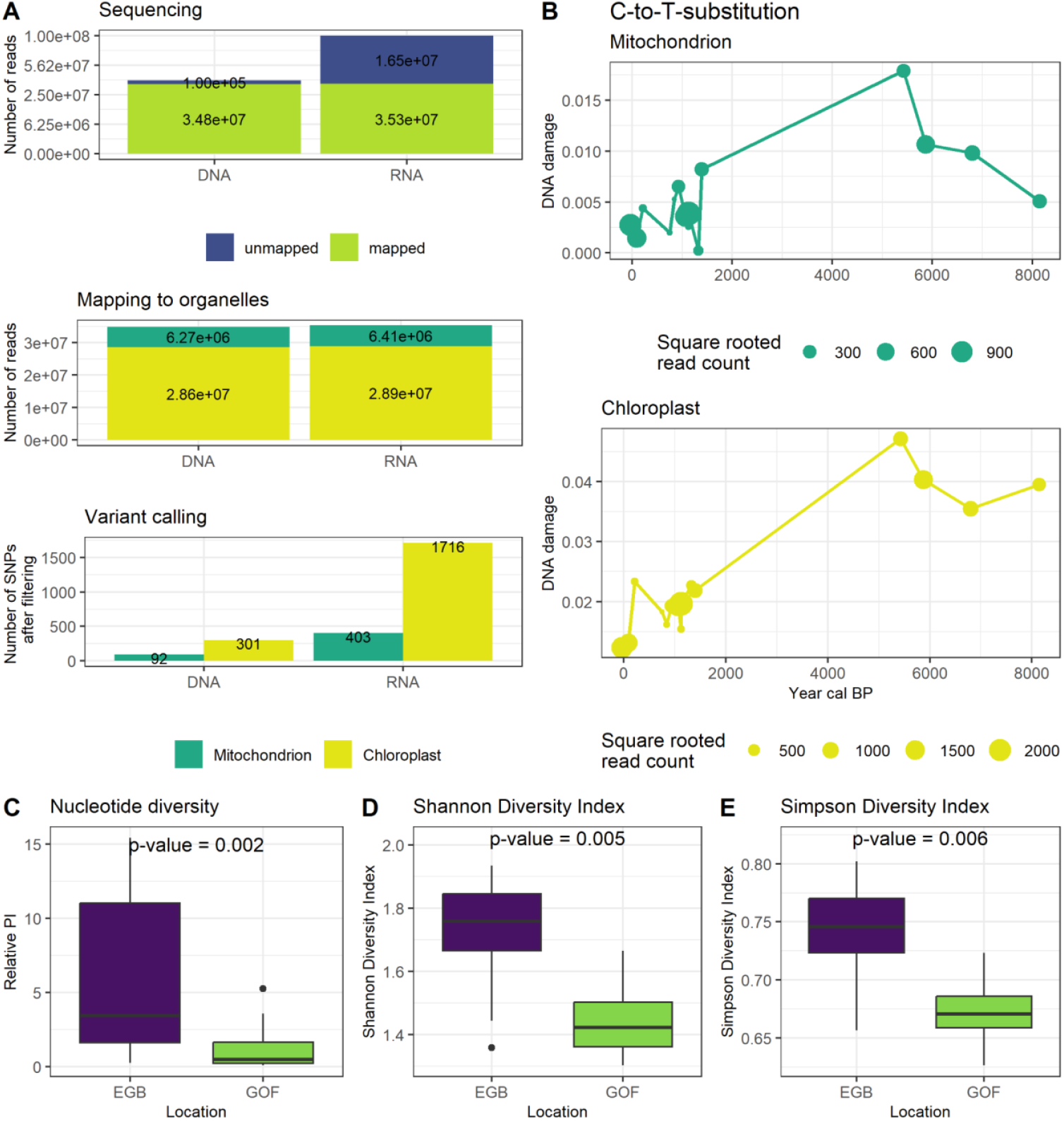
Data assessment: A) Sequencing Yield and Single Nucleotide Polymorphisms (SNPs): Post-trimming sequencing yield for DNA and RNA baits, and the number of SNPs detected in mitochondrial and chloroplast genomes for each bait set. B) C-to-T Substitution: Analysis of C-to-T substitutions over time for each organelle from all data (GOF & EGB). C-E) Comparative analysis of normalized nucleotide diversity and diversity indices. C) Relative Nucleotide Diversity (PI), D) Shannon, and E) Simpson between Eastern Gotland Basin (EGB) and Gulf of Finland (GOF). Each bar graph shows data for both sites with significant p-values.

The average coverage achieved for the experimental samples was generally high, with an average mitochondrial coverage of 78.77 and an average chloroplast coverage of 575.5. This coverage demonstrates the efficiency of the hybridisation capture approach and ensures reliability of the genomic data collected. Furthermore, the Extraction Blanks and Library Blanks produced low to non-meaningful coverage (see Supplementary Table S2). All information about the samples, corresponding age, location and other metadata is available in Supplementary Table S3.

### Ancient DNA – damage pattern

We analyzed damage patterns in ancient DNA (aDNA) samples, focusing on C-to-T substitutions. These substitutions are key aDNA authentication markers due to their pervasiveness in aDNA and their function in indicating cytosine deamination, a prevalent form of DNA damage. No significant correlation (R = 0.18, p = 0.315) was found between mapping coverage and substitution frequency, suggesting that damage pattern does not affect the mapping coverage. The degree of damage was found to significantly increase with the age of the samples (R = 0.65, p = 0.0003, Fig. 2B). A greater number of sequences were mapped to the chloroplast due to its larger genome size. This allows a greater number of reads to be used to detect damage patterns, increasing the overall value of the data.

### Genetic diversity and spatial differentiation

A comparison of the genetic differences between *S. marinoi* retrieved from the EGB and the GOF respectively, revealed that while the two locations exhibited some differences, they also shared a considerable degree of similarity. This was evidenced by the FST value, a measure used in population genetics to quantify the genetic differentiation between populations, which ranged between 0.05 and 0.1 (see Supplementary Material, Fig. S6). Notably, the genomic data from EGB shows a higher level of genetic diversity than the one from GOF. This is presented by different measures of distinct variant diversity, with significant differences observed (p-values: 0.002, 0.005, 0.006, see Fig. 2 C-E).

### Effects of environmental change

Our study on *S. marinoi* organelles at two locations in the Baltic Sea reveals that across several millennia the genetic composition of the population remained stable, punctuated by differences in specific periods, putatively coinciding with environmental changes. A Principal Component Analysis (PCA), conducted to investigate differences in allelic composition between sediment layers, showed that the majority of samples are situated within a primary cluster, but a number of smaller clusters are found that correspond with periods marked by specific environmental events (Fig. 3A). Together, PC1 and PC2 describe 42.6% of the data variance.

**Fig. 3:**
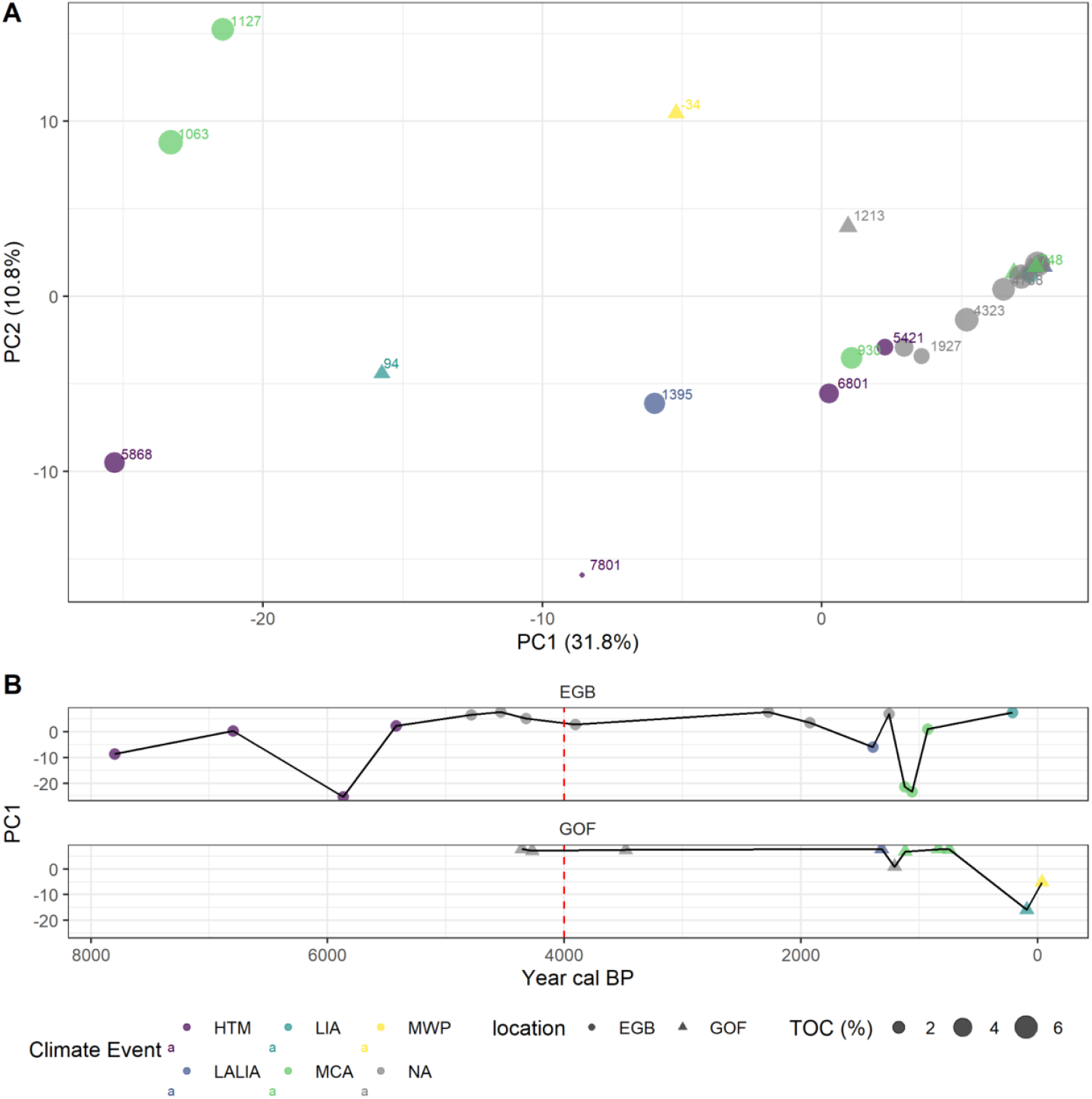
Principal component analyses of the allelic composition of *Skeletonema marinoi* organelles, demonstrating the genetic diversity and population structure. A) PC1 and PC2 are categorized by climate events, total organic carbon (TOC) and the age of each sample is given for both sites. B) PC1 versus time for both sites. The red dashed line marks the transition from the Littorina Sea to the Modern Baltic Sea. Climate events include the Holocene Thermal Maximum (HTM), Late Antique Little Ice Age (LALIA), Medieval Climate Anomaly (MCA), Little Ice Age (LIA), Modern Warm Period (MWP) and NA = No event

The retrieved specific clusters (Fig. 3A) correspond to specific climate periods, including the Holocene Thermal Maximum (HTM, 10,000-6,000 cal yr BP), Little Ice Age/Late Antique Little Ice Age (LIA, 550-100 cal yr BP/LALIA, 1,414-1,290 cal yr BP), Medieval Climate Anomaly (MCA), and Modern Warm Period (MWP). We verified that existing variation in allelic composition is not a result of the C-to-T-substitution rate by performing a PERMANOVA, which showed that the C-to-T-substitution rate accounted for approximately 6.7% of the variation in the PCA space (R^2^ = 0.067, p = 0.19), though this was not statistically significant (see Supplement, Fig. S5). Additionally, we performed a PERMANOVA to assess the impact of Total Organic Carbon (TOC) on the variation in the PCA space, which showed that TOC accounted for approximately 4.5% of the variation (R^2^ = 0.045, p = 0.328), also not statistically significant.

In the EGB, the period under consideration extends to the Ancylus Lake (extrapolated age: 8,148 cal yr BP), and responses are observed during the HTM, LALIA, MCA, and LIA periods. The GOF record, which covers the last ca. 4,300 years, displays changes during the LIA and MWP periods. During periods without profound climate events, we observe a consistency in allelic composition, which points toresilience of the algal populations over time. However, within warm periods (e.g. HTM and MCA), there is a notable degree of variability, particularly for EGB. This pattern of long-term stability and resilience, interrupted by distinct variation, is visualized in Fig. 3B, depicting changes along PC1 in both cores. This shows temporal variations that align with specific climate events, suggesting that the population’s genetic makeup is undergoing changes in response to these events. Additionally, nucleotide diversity increased significantly during the transition from the Littorina Sea to the Modern Baltic Sea, around 4,000 cal yr BP, and then successively decreased again (Supplementary Material, Fig S7).

Allele turnover, a measure of the rate at which new alleles replace old ones, provides insights into genetic variation over time. Figure 4 displays patterns of genetic change over time. A Generalized Additive Model (GAM) was used to analyze allele turnover as a function of time.

**Fig. 4:**
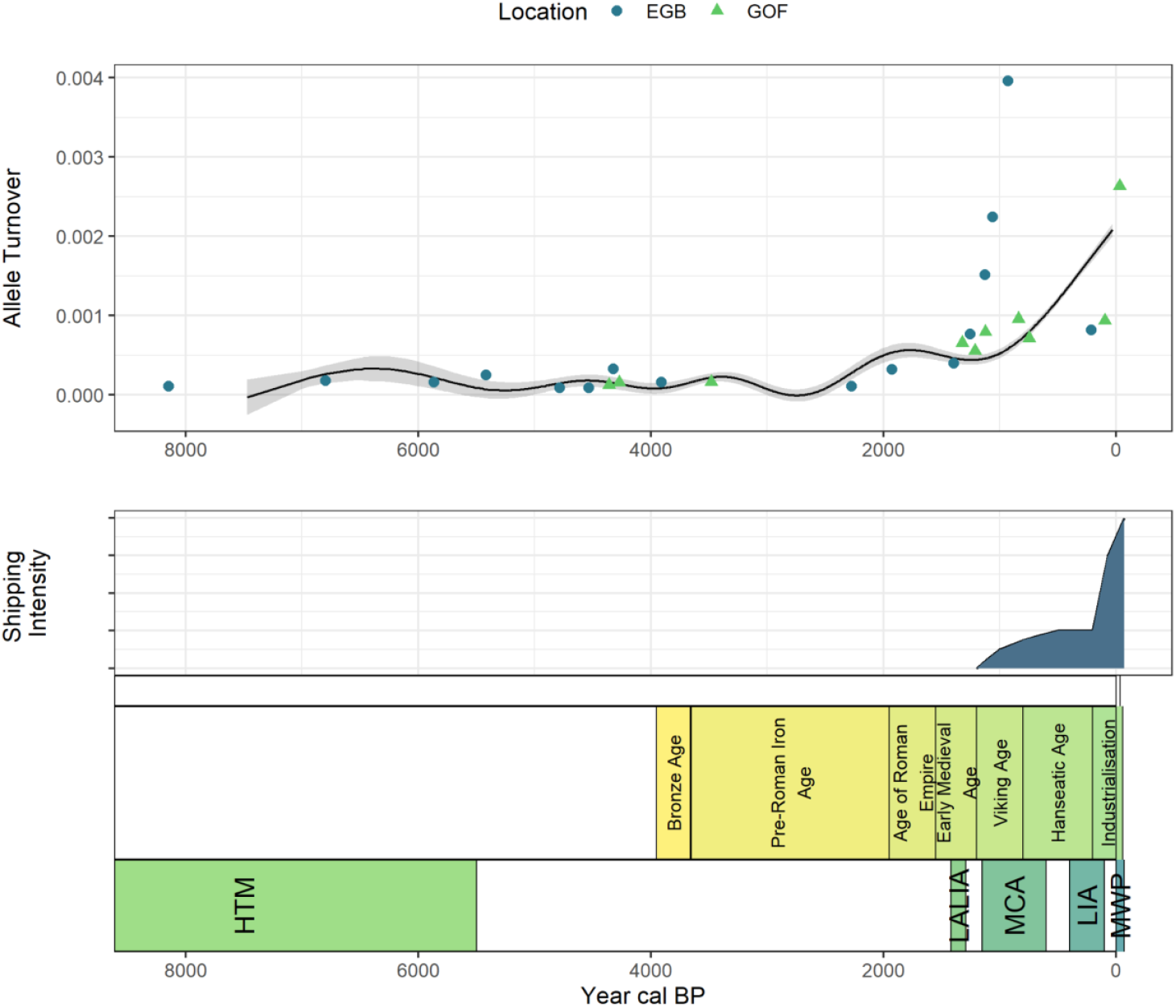
Temporal Analysis of Allele Turnover in *Skeletonema marinoi*. This figure presents the Generalized Additive Model (GAM) fitted to the allele turnover over time (BP). The data includes both Eastern Gotland Basin (EGB) and Gulf of Finland (GOF) sites. Key environmental events, such as the Holocene Thermal Maximum (HTM), Late Antique Little Ice Age (LALIA), Medieval Climate Anomaly, Little Ice Age (LIA), and Modern Warm Period (MWP), are noted. Additionally, anthropogenic periods and associated shipping activity estimated from literature sources are shown.

At EGB, an increase in allele turnover is observed around 1,500-1,000 cal yr BP, indicating a potential for heightened genetic change from the Medieval period onwards. At GOF, turnover remains consistent until around 94 to -34 cal yr BP. While industrialization and anthropogenic impacts may be associated with recent genetic shifts, establishing a causal link between these changes and earlier historical periods requires caution.

## Discussion

The millennial-long stability, spanning from the Ancylus Lake (approx. 8,148 calibrated years BP), to Littorina Sea and Modern Baltic Sea to 1,500–1,000 cal yr BP in the Eastern Gotland Basin (EGB), and from around 4300 to 94–34 cal yr BP in the Gulf of Finland (GOF), aligns with findings from other phytoplankton studies, albeit with a different temporal perspective. The findings of these studies indicate that long-term genetic stability is frequently maintained in marine populations due to their substantial population size and high dispersal capacity, even in the face of significant environmental changes ^16^. These factors contribute to a buffering effect against genetic drift and local extinctions, allowing populations to maintain genetic diversity over long periods of time. The ability of *S. marinoi* to cope with changing environmental conditions through reversible shifts in genetic composition as shown here for the Baltic Sea further supports this stability. This resilience is a common trait among phytoplankton, allowing them to respond to environmental changes without losing overall genetic diversity^17^. The observed stability is expected given the ecological and evolutionary characteristics of phytoplankton populations. Large population sizes, high reproductive rates, and the potential for gene flow between populations contribute to the maintenance of genetic diversity and stability over long periods of time^18^. Other findings, such as studies of diatoms and other phytoplankton species have reported patterns of genetic stability and adaptability of phytoplankton to environmental change^19^. These findings imply that the observed stability in *S. marinoi* populations is not only expected, but also indicative of the broader ecological and evolutionary dynamics of marine phytoplankton. Our data demonstrates that this stability can be upheld across several millennia.

However, this stability is not without punctuated interruptions. Our results show that climatic events have an influence on the allelic composition of *S. marinoi* populations. For example, shifts in the genetic composition of the EGB population were observed during the Holocene Thermal Maximum, the Late Antique Little Ice Age, and the Medieval Climate Anomaly. Similarly, the GOF population showed changes during the Little Ice Age and the Modern Warm Period. These shifts highlight the ability of *S. marinoi* to cope with different climatic events. The discrepancy between EGB and GOF may be due to the slightly higher genetic diversity in the EGB (Fig. 2C-E), which may facilitate adaptation. The differences between the two sites can also be attributed to their distinct geographical characteristics. The EGB, located in the Baltic Proper, may experience an influx of migrants from other populations, increasing genetic diversity and resilience. In contrast, the genetic composition of the GOF population could be influenced by its unique environment, characterized by freshwater influx and lower salinity levels^20^, as well as more extensive ice cover in the northern regions of the Baltic Sea^21^. In particular, samples from the Little Ice Age and the Modern Warm Period show more pronounced responses in the GOF. The allelic composition remains relatively constant in the absence of climatic events, indicating a stable genetic structure under stable conditions and once a shift has manifested.

After this long period of punctuated genetic stability, recent centuries reveal a novel pattern of rapid allelic turnover, and this coincides with increased human activity. This rapid turnover is particularly evident in the GOF during periods such as the pre-Roman Iron Age, the Viking Age, the Hanseatic Age, and the Industrial Revolution. These results highlight the possible impact of human activities on the genetic dynamics of *S. marinoi* populations. This increased activity is manifested by increased shipping activity in the Baltic Sea^22,23^. Ballast water exchange^24^ in the last two centuries could have been a specific agent of increased changes in haplotype composition, causing rapid translocation of populations. This rapid allelic turnover contrasts with the more gradual and reversible shifts observed in response to natural climatic events, highlighting the influence of anthropogenic factors on the genetic composition of these populations. Due to its coastal position, the GOF is more exposed to these changes, and perhaps the phytoplankton is also more directly affected.

The rapid allelic turnover observed in recent centuries, particularly in response to human activities, underscores the influence that anthropogenic factors may have on the genetic dynamics of populations. Our study on the population dynamics of *S. marinoi* populations provides valuable insights into the long-term resilience and adaptability of marine phytoplankton. The observed genetic stability over several millennia, punctuated by reversible shifts in response to climatic events, underscores the inherent resilience of these populations. However, the recent rapid allelic turnover highlights the potential impact of human activities on population dynamics and structure, and thus presents an opportunity for further investigation.

At the same time it is important to acknowledge the limitations of our study. The limited number of samples included may affect the overall strength and generalizability of our findings. In addition, the lack of precise tests limits our ability to draw definitive conclusions about long-term genetic trends and the full extent of human impact. To date, we also lack a sufficient understanding of the distribution and taphonomy of ancient eDNA, but in sediments, DNA of small organisms, such as phytoplankton, has been shown to give a representative signal of a water body^25^. The data of the two cores display a high level of congruency, and despite the limitations, this indicates a link between environmental change, human activity, and genetic dynamics. Future studies with larger sample sizes and e.g. more locations are needed to further explore the complex interactions between environmental change and genetic responses.

The results of our study provide a solid base for future research on the effect of current global change on the genetic composition of natural populations across millennia. By understanding the genetic stability and adaptability of *S. marinoi* populations, we can better understand ecological and evolutionary dynamics of marine phytoplankton. This knowledge is crucial for predicting how these populations may respond to current and future environmental changes, including climate change and direct anthropogenic pressures, and thus offer a valuable foundation for future conservation efforts. Our study provides novel insights into the genetic resilience and adaptability of *S. marinoi* populations. It demonstrates that these populations maintain stability over millennia and respond dynamically to both natural climate fluctuations and human-induced changes. In sum, it highlights the importance of considering both natural and anthropogenic drivers and their relative impacts when assessing and predicting genetic resilience of marine ecosystems.

## Material & Methods

### Study area

The Baltic Sea is a relatively young and shallow brackish water system currently facing a significant risk of increasing hypoxia, a condition characterized by low oxygen levels and bottom water anoxia. This phenomenon is primarily attributable to the reduction in dissolved oxygen and nutrient discharge resulting from warmer water temperatures, which in turn promote the formation of algal blooms^26^. These blooms decompose and consume oxygen, leading to the development of hypoxia. In recent history, the Baltic Sea has experienced a notable increase in eutrophication, a process driven by the excessive input of nutrients such as nitrogen and phosphorus from agricultural runoff, wastewater discharge, and industrial activities^13^. The enrichment of nutrients has resulted in the proliferation of phytoplankton and algal blooms, which, upon decomposition, have further depleted oxygen levels in the water. The Baltic’s unique position as a land-enclosed sea with high human and industrial activity, coupled with its low biodiversity, makes it particularly vulnerable to these changes.

The Baltic Sea’s history is well known. It has undergone substantial environmental changes in the past. After the end of the last glaciation approximately 10,000 years ago, it went through several fresh- and saltwater stages, starting as a large meltwater lake, then transitioning into various states of salinity due to glacial retreat and land rebound^27^. These stages included the Baltic Ice Lake, the Yoldia Sea, the Ancylus Lake, and the Littorina Sea^28^. The Baltic Sea’s temperature and salinity have fluctuated over time, with notable increases during the Holocene Thermal Maximum and the Medieval Climate Anomaly, and a recent increase since 1850^27,29,30^.

Within the Baltic Sea this study covers a time series of two distinct locations: The Eastern Gotland Basin, located in the Baltic Proper with a profound water depth of max. 249 meters, and the comparatively shallower Gulf of Finland, reaching a depth of 81 meters. At both locations, we collected sediment cores to inform on the effects of environmental changes on the population genetic composition of *S. marinoi*.

### Sampling

Two sediment cores were taken (Fig. 1A) in April 2021, during expedition EMB262. 1) Eastern Gotland Basin (EGB, 57°17.004’N, 020°07.244’E, 241 m water depth, core EMB262/6-30GC); 2) Gulf of Finland (GOF, 59°34.443’N, 023°36.461’E, 81 m water depth, core EMB262/12-3GC). The cores (Fig. 1B, ca. 500 cm) were taken using a gravity corer (GC). The sampling for the EGB began at a depth of 32 cm. The cores were sampled on board of the research vessel using sterile syringes according to Epp et al.^31^ and immediately frozen for storage.

Core dating was performed as described in Schmidt et al.^32^. This was done by correlating organic carbon records from post-Littorina transgression sediments in the Baltic Sea. The chronology of the cores is determined by the relative S content and the Br/K ratio. The Br/K ratio reflects changes in the bulk organic carbon content of the sediments. The data are visually matched with XRF and organic carbon data from dated sediment cores from the same or nearby locations for different historical periods. For the sediment core from the Gulf of Finland, an independent Bayesian age model was constructed based on radiocarbon dates of bulk organic matter in the sediments. This allows the sediments to be accurately dated to around 2300 BC. The same core samples were used as in Schmidt et al.^32^ and in this study.

## Laboratory work

### DNA extraction

The PowerSoil Pro Kit from Qiagen (Hilden, Germany) was used to extract DNA from both sediment cores (n=26, see supplementary Table SX) covering various depths (ages). For the extraction 0.5 g of sediment was used per sample, resulting in 26 samples and 2 extraction blanks. The extraction process was carried out according to the manufacturer’s protocol with some modifications including an overnight incubation at 56 °C with the addition of 20 μL proteinase K (20 mg/ml) to enhance sample lysis. The washing steps were performed using a Qiagen Vacuum Pump (Hilden, Germany). Subsequently, samples were centrifuged for 3 minutes. After centrifugation, the elution process was performed in two steps. Each part involved adding 75 μL of elution buffer to the sample, allowing it to incubate for 5 minutes, and then collecting the eluate. Both eluates were collected in the same tube.

### Library preparation

A single-stranded library approach according to Gansauge et al.^33^ was performed to maximize the retrieval of genetic information from these degraded sedaDNA samples. This method, optimized for fragmented DNA, has been shown to significantly increase the sequence yield and preserves unique molecules, thereby providing a more comprehensive genomic analysis. The library preparation was conducted with 40 ng input DNA. All 26 samples and 2 extraction blanks were processed in the library preparation. Additionally, 2 library blanks were created.

After the indexing PCR, each library was cleaned using the MinElute PCR purification Kit (50,250; Qiagen) resulting in a final amount of 20 μL per sample (For more detailed info, see Supplement Section 1.1.). The samples were then analyzed on a 2100 Bioanalyzer (Agilent Technologies, Santa Clara, CA, USA) using the Agilent Bioanalyzer High Sensitivity DNA Analysis Kit (Agilent Technologies, Santa Clara, CA, USA). Due to adapters still being present in the samples, we performed an additional purification using the HighPrep™ bead cleanup (MagBio Genomics Inc., Gaithersburg, MD, USA). 1.6x of HighPrep™ PCR reagent was used. The DNA concentration was measured using ds-DNA HS Assay Kit and the Qubit® 4 fluorometer (Invitrogen Thermo Fisher, Waltham, MA, USA). The libraries were combined into pools of four, each with the same DNA amount. The extraction and library blanks were incorporated in the same volume as the sample with the lowest concentration. The libraries were then further used for hybridization enrichment of chloroplast and mitochondrial genomes.

### Hybridization enrichment

In this study, a hybridization enrichment approach was employed. Hybridization enrichment allows for targeted isolation of specific sequences, improving sequencing efficiency by reducing the presence of non-target DNA. Thus, providing a better representation of the studied species, enabling a more detailed and precise analysis of its genetic composition.

### Bait design

The design of the bait set was carried out for two target reference organelle genomes: the mitochondrion and the chloroplast (GenBank PRJNA493755). Together, these genomes consist of 170,788 nucleotides (nt), with 127,202 nt from the Chloroplast and 43,586 nt from the mitochondrion, and an average GC content of 30.9%. To improve the accuracy of the baits, the contigs were softmasked for simple and low-complexity repeats.

Subsequently, baits were designed with a length of 80 nt and a tiling coverage of 4x. This means that, on average, each nucleotide in the target sequence is covered by four different baits, thereby ensuring redundancy and enhancing the probability of successful hybridization. This process led to the generation of 10,000 probes. The baits were then filtered based on softmasking for simple and low complexity repeats. Only those with 35% or less softmasking retained. A total of 7,985 baits passed this filtering step.

### Enrichment

This bait design was then used for two different bait sets: 1) RNA baits (BioCat, Heidelberg, Germany); 2) DNA baits (Integrated DNA Technologies, IDT). The libraries were enriched with both RNA and DNA baits, resulting in two sets of libraries: 1) RNA baits enriched; 2) DNA baits enriched. General steps are explained in the following. More detailed information on the used protocols can be found in the supplementary data (section 1.2.).

The enrichment with RNA baits was conducted according to the myBaits v.5.02 manual. A two-round enrichment strategy was performed. The hybridization mix was prepared and transferred to a rotation oven once the 24-hour incubation at a hybridization temperature of 63°C started. In the following bead cleanup, the libraries were resuspended and amplified. Each 27.4 μL PCR reaction included the following nine components: 1) 3.45 μL of DEPC treated H_2_O; 2) 2,5 μL 10X HiFi PCR Buffer (Invitrogen Thermo Fisher, Waltham, MA, USA); 3) 0.25 dNTPs (25mM); 4) 1 μL BSA (20 mg/ml); 5) 1 μL MgSO_4_ (50 mM); 6) 0.2 μL Platinum™ Taq DNA-Polymerase High Fidelity (5 U/μL) (Invitrogen Thermo Fisher, Waltham, MA, USA); 7) 2 μL IS5_bridge_P5 (10 μM); 8) 2 μL IS5_bridge_P5 (10 μM), 9) 15 μL library pool. The amplification was run using the following settings: 1 minute initiation at 94°C, 14 cycles of 15 seconds denaturation at 94°C, 20 seconds annealing at 60°C and 1 minute extension at 68°C, followed by 2 minutes of final elongation at 68°C and afterward the sample was stored at a temperature of 20°C. Afterwards, the samples were purified with beads and eluted in 10 μL. The second round of enrichment was performed in a similar way. The PCR was conducted in 8 cycles and the pools were eluted in 17 μL after the bead purification (see more detailed in supplement section 1.2.2.).

Enrichment with DNA baits was also performed in two rounds following the IDT (Integrated DNA Technologies, Löwen, Belgium) xGen™ hybridization capture of DNA libraries protocol (option: AMPure XP Bead DNA concentration protocol), with the following modification. The hybridization temperature was set to 63°C. Once this temperature was reached on the thermal cycler, the tubes were moved to a rotation oven and incubated for 16 hours (see more detailed in supplement section 1.2.1.).

All resulting pools were quantified using Qubit® ds-DNA HS Assay Kit and Agilent Bioanalyzer HS Kit. Afterwards, the pools enriched with RNA baits were collectively combined into a single pool, while those enriched with DNA baits were combined into another separate pool, each in equal DNA amounts. This resulted in two distinct library pools. Both of these pools were sent for Illumina sequencing (paired-end, 2 × 150 bp, MiSeq-v3) at Fasteris SA (Geneva, Switzerland).

### Bioinformatics

The sequences were trimmed, mapped against references, and converted to finally calculate mapping coverage and for variant calling. First, sequence file names were modified to fit the sample identifier using python v3.10.13. Then raw sequence reads were processed using Autotrim v0.6.1, a tool that automates the quality control and trimming of high-throughput sequence data^34^. Autotrim employs Trimmomatic v0.39^35^, to trim sequences based on quality and remove adapters, FastQC v0.12.1^36^ to assess the quality of the raw sequence data, and MultiQC v1.14 ^37^ to aggregate the results into a single report. Following the initial processing, the trimmed reads were mapped to the reference chloroplast and mitochondrial genomes (GenBank: PRJNA493755) using BWA-MEM v0.7.17^38^. BWA is a software package for mapping low-divergent sequences against a reference genome, which is particularly useful for precisely aligning sequencing reads.

The mapped reads were then converted to bam format using Samtools v1.15^39^, a suite of programs for interacting with high-throughput sequencing data. In addition, the per-sample mapping coverage was calculated. After careful consideration, we decided against deduplicating our sequence data as deduplication resulted in a significant loss of data. This is probably due to the nature of our short and fragmented sequences. In addition, each variation, even those present in duplicates, may hold biologically significant information. Therefore, removing duplicates may have eliminated critical genetic diversity, compromising the integrity of our analysis.

VCFtools v0.1.16^40^ and BCFtools v1.19^39^ were used for calling and filtering variants from the alignment. Variant calling is the process of identifying differences, such as single-nucleotide polymorphisms (SNPs) and insertions/deletions (indels), between the sequenced sample and the reference genome. Variants were filtered to include only those with a quality score (Q) of 30 or higher. The identified variants were then phased into haplotypes using Beagle v5.4^41^. Haplotyping, or phasing, is the process of determining the distribution of alleles of multiple variants along each chromosome. In this context, it can be said it accounts for unique combinations of alleles at variant sites (distinct variations) across the chloroplast and mitochondrial genome.

The pattern of ancient DNA damage was assessed using mapDamage2 v2.2.2^42^. This tool quantifies patterns of DNA damage in ancient samples, which can provide insights into the preservation and authenticity of ancient DNA.

### Analyses

The bioinformatic output was converted into a table format and for further analyses with R v4.3.1: The mapping coverage (i.e., the number of reads mapped against reference genome), allele frequencies, distinct variants, nucleotide diversity and C-to-T-substitution rate were combined with the sample metadata (Supplement, Tab. S3) using the R package “tidyverse”^43^. The data was visualized using the packages “tidyverse’”, “ggpubr” v0.6.0^44^ and “tidypaleo” v0.1.3^45^.

In order to facilitate a comparison of the genetic diversity between GOF and EGB, nucleotide diversity (resulting from Samtools) and distinct variants (resulting from Beagle) were normalized based on the mapping coverage for mitochondria and plastid. Second, Pearson’s correlation was employed to verify the consistency of nucleotide diversity between the mitochondrion and chloroplast. For comparison, the dataset was subset to the overlapping time period of both locations. Subsequently, a t-test was applied to compare and test the significance of the nucleotide diversity (mitochondrial and chloroplast data combined) between the two sites. In addition, each distinct genetic variant was assigned a unique identifier for easy reference. These identifiers were utilized to calculate two diversity indices: the Shannon index and the Simpson index. These indices were compared between the EGB and GOF sites using a t-test,

To further identify spatial differences between GOF and EGB, we analyzed allele frequencies (resulting from VCFtools & BCFtools) and calculated the Fixation Index (FST) using corresponding samples from both locations. The FST is a measure of population differentiation attributable to genetic structure. The FST value ranges from 0 to 1, with 0 indicating complete interbreeding (i.e., the two populations are freely intermixing) and 1 indicating that all genetic variation is explained by the population structure (i.e., the two populations do not share any genetic diversity). We implemented a custom R function to calculate the FST over time. This process involved sorting the data by age, subsetting samples within a specified time window, and computing FST based on allele frequency variance. The allelic composition, derived from parsing VCFtools output, was analyzed using Principal Component Analysis (PCA) to examine the genetic structure.. We also performed Pearson’s correlation and PERMANOVA to assess the impact of C-to-T substitution rates and Total Organic Carbon (TOC) on the variation in the PCA space, specifically the first two principal components (PCs). These analyses helped determine the extent to which these factors influenced genetic composition. For PCA and PERMANOVA we also used the functions provided by the package “vegan” v2.6.4^46^.

The C-to-T-substitutions at the first position from the mapDamage2 output were examined to authenticate the ancient DNA (aDNA) and analyze the damage patterns. A Pearson’s correlation test was conducted to assess the influence of mapping coverage on these substitutions. This was done to evaluate if coverage has an impact on the observed damage. The Baltic Sea Mn/Ti ratio data were also integrated into the analysis.

To assess the genetic diversity and adaptability of the population, we calculated the rate of allele turnover. This measure, derived from the ratio of significant allele changes (greater than 1%) to total generations, reflects the estimated rate of genetic change over time based on the present data. Given the low FST values, we inferred that the two sites, EGB and GOF, belong to a single population. The calculation covered all of the data, from the oldest sample from EGB to the youngest sample from GOF, providing insight into the evolving genetic makeup of the population.

To identify phases of change in the allele turnover across the entire dataset, a generalized additive model (GAM) was fitted using the “gam” function from the “mgcv” package^47^. This model was applied to the complete dataset, including periods where we did not have direct samples, as the allele turnover was estimated based on the available data. We calculated the mean allele turnover for the present data points, which were then included in the GAM plot. Allele turnover was used as a response variable to fit a gam model as a function of “age”.

Corresponding metadata to this analysis can be found in supplementary material (Table S3). Data on shipping activity in the Baltic Sea were obtained from three main sources: Kontny^48^, Mägi^49^, and Rytkönen et al.^50^. Kontny’s work provided insights into maritime contacts during the Roman and Migration Periods (1st-7th centuries AD), while Mägi’s research focused on the role of the Eastern Baltic in Viking Age communication across the Baltic Sea. Rytkönen et al.’s research on the statistical analysis of Baltic Sea shipping revealed a significant increase in maritime traffic during the late 20th and early 21st centuries. These sources were used to establish a timeline of shipping intensity from 1200 BP to the present.

## Acknowledgements

This study was funded by the K314/2020 grant of the Collaborative Excellence Programme of the Leibniz Association. The authors acknowledge support by the High Performance and Cloud Computing Group at the Zentrum für Datenverarbeitung of the University of Tübingen, the state of Baden-Württemberg through bwHPC and the German Research Foundation (DFG) through grant no INST 37/935-1 FUGG. AS acknowledges funding from the International Max Planck Research School for Quantitative Behaviour, Ecology and Evolution. MB is supported by the Hessian Ministry of Higher Education, Research and the Arts through the LOEWE Centre for Translational Biodiversity Genomics. For help with preparing reagents for library preparation and enrichment experiments we thank Clara Linde Schlimbach and Anna Weis. For setting up the ancient DNA clean laboratory in Frankfurt where all DNA extractions were done by AS and JR, we thank Leonie Schardt. The authors declare no conflicts of interest.

## AI statement

In the preparation of this manuscript, we employed artificial intelligence (AI) tools to enhance the quality of our work and to optimize our research process.

R Script Optimization: We utilized an AI-based code optimization to refine our R scripts. The tool provided suggestions for code enhancement, identified potential bugs, and recommended more efficient coding practices. This not only improved the performance of our scripts but also ensured the reproducibility and reliability of our results.

We believe that the use of AI in our research process has improved the quality of our work. However, we stress that the final decisions on manuscript content and the interpretation of the results were made by the authors.

## Data availability

The datasets generated and analyzed during this study are publicly available. Data and scripts can be accessed through the GitHub repository at https://github.com/Alex-132/seda_enrichment.

Additionally, sequence data have been deposited in the European Nucleotide Archive (ENA) under the accession number XXXXX (will be uploaded there).

