## Supplement for "Multi-millennial genetic resilience of Baltic diatom populations disturbed in the past centuries"

#### 1. Supplementary Materials

##### 1.1. Single stranded library preparation

The used library preparation protocol conducted a single stranded approach and was performed according to Gansauge et al. (2020) publication. The steps are detailed in the following section:

###### Day 1 - Adapter Ligation

**Before Starting:**

- **Aliquot the reagents** before beginning the experiment.
- **Prepare 2% Tween-20** by combining 98 µL of DEPC water with 2 µL of Tween-20.
- **Prepare a 0.1 pM dilution for the CL304** (spike-in) to use as a positive control by mixing 1 µL of the 10 pM solution with 99 µL of TET buffer.
- **UV Treatment:** Prepare the following items:
    - 2 x PCR strips
    - 2 x 1.5 mL tubes
    - DEPC water aliquot
    - 2 x 1.5 mL TET buffer aliquot

**Equipment Required:**

- Thermocycler
- 0.2 mL tubes
- Cooling rack for 0.2 mL tubes

**Preparation:**

- Prepare both mastermixes in advance.
- Work quickly to maintain sample integrity.

**Oligonucleotide-Dilution:**
Before starting the library preparation experiment:

1. Thaw one tube of the spike-in positive control (CL304) at room temperature.
2. Vortex the tube and spin down the liquid.
3. Prepare a 0.5 nM dilution by combining:

- 5 µl of the diluted oligonucleotide with
- 995 µl of TET.

4. Vortex the tube and spin down the liquid again.
5. Prepare a final 10 pM oligonucleotide dilution by combining:
   - 2 µl of the previous dilution with
   - 98 µl of TET.
6. Vortex the tube and spin down the liquid.

**1. Heat Denaturation & Dephosphorylation**

**Reagents for Reaction Mix:** (Total Volume: 45.6 µl)

| **Reagent** | **Volume (µl)** | **Total Volume (µl)** |
|---|---|---|
| Mastermix | 17 | 17 |
| 10x T4 RNA ligation buffer | 8 | 136 |
| 2% Tween-20 | 2 | 34 |
| Water (HPLC grade) | 4.6 | 78.2 |
| Spike-in positive control CL304 (10 pM) | - | - |
| Fast AP (1 U/µl) | 1 | 17 |
| **Total Reaction Mix** | **15.6** | **282.2** |
| DNA sample | 30 | |
| **Total Volume** | **45.6** | |

**Procedure:**

1. Mix the components and spin down in a microcentrifuge.
2. Incubate the reaction in a thermocycler for 10 minutes at 37°C.
3. Heat the mixture to 95°C for 2 minutes, then immediately place it on a cooling rack.

**2. Ligation of the First Adapter**

**Reagents for Ligation Reaction:** (Total Volume: 80 µl)

| **Reagent** | **Volume (µl)** | **Total Volume (µl)** |
|---|---|---|
| Mastermix | 17 | 17 |

| Reagent | Volume (µl) | Total Volume (µl) |
|---|---|---|
| 50% PEG-8000 (NEB) | 32 | 544 |
| ATP (100 mM, ThermoFisher) | 0.4 | 6.8 |
| TL181/TL159 (10/20 µM) | 1 | 17 |
| T4 DNA Ligase (30 U/µl) | 1 | 17 |
| **Total** | **34.4** | **584.8** |
| Add Mixture from Step 1 | 45.6 | |
| **Total Volume** | **80** | |

**Procedure:**

1. Pre-mix the reagents in a 2 ml tube by repeatedly inverting the tube on a rotator for 5 minutes before adding the enzyme.
2. Invert the tubes upside down on a rotator for at least 10 additional minutes to ensure proper mixing.
    - **Note:** PEG-8000 is highly viscous; pipette slowly. Vortexing is not recommended as it does not achieve homogenous mixing.
3. Mix and spin down in a microcentrifuge. Visually inspect each tube to ensure there are no streaks.
4. Incubate the ligation reaction at 37°C in a thermocycler for 1 hour.
5. Incubate for an additional 2 minutes at 95°C in the thermocycler, then hold at 10°C.
6. Freeze ligations at -20°C.
7. **Break:** Take a break before proceeding to the next steps.

#### Day 2 - Library Preparation

**Equipment Required:**

- 1.5 mL tubes
- Thermomix (1.5 mL, set to 45°C and 25°C)
- Thermocycler (95°C)
- Cooling rack for 0.2 mL tubes
- Magnetic rack for 1.5 mL tubes

**3. MyOne Bead Preparation**

1. **Resuspend Dynabeads MyOne Streptavidin C1** by vortexing in 2 mL LoBind tubes.

2. In a separate 2 mL tube, resuspend:
    - **320 µL (16 x 20 µL) of beads** in **500 µL Binding Buffer I (B&W buffer I)**.

| **Reagent** | **Volume (µL)** |
|---|---|
| Mastermix | 16 |
| MyOne C1 beads (x16 Samples) | 320 |
| B&W Binding Buffer I | 500 |

3. Resuspend by vortexing and spin the tubes in a microcentrifuge.
4. Pellet the beads using the magnetic rack (place on the magnetic rack).
5. Remove the supernatant.
6. **Second Wash:**
    - Add **500 µL B&W Buffer I**.
    - Resuspend by vortexing and spin the tubes in a microcentrifuge.
    - Pellet the beads with the magnetic rack and remove the supernatant.
7. Resuspend the beads with **B&W Buffer I**. Use **100 µL for each reaction + 100 µL extra (Master Mix)**.
8. Aliquot the suspension into **100 µL** in 1.5 mL tubes.
9. Add **100 µL B&W Buffer I** to the ligation reaction from Day 1.
10. Transfer the diluted ligation reaction into the bead suspension and mix by pipetting.
11. The final volume should be **280 µL**.
12. Invert the tubes upside down on a rotator for **20 minutes at room temperature**.

**4. First Bead Wash**

1. Spin the beads briefly.
2. Pellet the beads using the magnetic rack and discard the supernatant.
3. Add **200 µL of B&W Buffer I** and resuspend the beads by vortexing. Spin the tubes briefly.
4. Pellet the beads with the magnetic rack and discard the supernatant.
5. Add **100 µL Stringency Buffer**.
6. Vortex and transfer to the thermomix set to **45°C**.
7. Incubate for **3 minutes** in the thermomix without shaking.
8. Pellet the beads with the magnetic rack and remove the supernatant.
9. Add **200 µL of Wash Buffer II** and resuspend the beads by vortexing.

10. Briefly spin the tubes and leave them in a rack in the hood.

**5. Primer Annealing and Extension**

**Prepare Fill-in Mix**

- Use 2 mL tubes.

| **Reagent** | **Volume (µL)** | **Total Volume (µL)** |
|---|---|---|
| Mastermix | 17 | 17 |
| DEPC Water | 41.6 | 707.2 |
| 10x Klenow Buffer | 5 | 85 |
| dNTPs (25 mM) | 0.4 | 6.8 |
| 2% Tween 20 | - | - |
| CL128 (100 µM) | 1 | 17 |
| Klenow Fragment (10 U/µL) | 2 | 34 |
| **Total Fill-in Mix** | **50** | **850** |

1. Use the bead suspension (200 µL from Step 4).
2. Spin down with a microcentrifuge.
3. Pellet the beads using the magnetic rack and remove the supernatant.
4. Add **50 µL of fill-in mix** and mix by pipetting.
5. Incubate the tubes at **35°C at 800 r.p.m.** for **20 minutes**.

**6. Second Bead Wash**

1. Spin the beads briefly.
2. Pellet the beads using the magnetic rack and discard the supernatant.
3. Add **200 µL of B&W Buffer I** and resuspend the beads by vortexing. Spin the tubes briefly.
4. Pellet the beads with the magnetic rack and discard the supernatant.
5. Add **100 µL Stringency Buffer**.
6. Vortex and transfer to the thermomix set to **45°C**.
7. Incubate for **3 minutes** in the thermomix without shaking.
8. Pellet the beads with the magnetic rack and remove the supernatant.
9. Add **200 µL of Wash Buffer II** and resuspend the beads by vortexing.

10. Briefly spin the tubes and leave them in a rack in the hood.

**7. Ligation of Second Primer and Library Elution**

**Prepare Ligation Mix**

- Use 2 mL tubes for mastermix.

| Reagent | Volume (µL) | Total Volume (µL) |
|---|---|---|
| Mastermix | 17 | 17 |
| DEPC Water | 76 | 1292 |
| T4 DNA Ligase Buffer (10x) | 10 | 170 |
| 50% PEG-4000 | 10 | 170 |
| CL53/TL178 (100 µM) | 2 | 34 |
| 2% Tween 20 | - | - |
| T4 DNA Ligase (5 U/µL) | 2 | 34 |
| **Total Ligation Mix** | **100** | **1700** |

1. Vortex the mixture before adding ligase.
2. Mix firmly by flicking the tube with a finger.
3. Spin down tubes with the bead solution (200 µL from Step 6).
4. Place the tubes on the magnetic rack and remove the supernatant.
5. Add **100 µL of ligation mix** and resuspend the beads by pipetting.
6. Incubate at **22°C for 1 hour**, shaking at **800 r.p.m.**.

**8. Third Bead Wash**

1. Spin the beads briefly.
2. Pellet the beads using the magnetic rack and discard the supernatant.
3. Add **200 µL of B&W Buffer I** and resuspend the beads by vortexing. Spin the tubes briefly.
4. Pellet the beads with the magnetic rack and discard the supernatant.
5. Add **100 µL Stringency Buffer**.
6. Vortex and transfer to the thermomix set to **45°C**.
7. Incubate for **3 minutes** in the thermomix without shaking.
8. Pellet the beads with the magnetic rack and remove the supernatant.

9. Add **200 µL of Wash Buffer II** and resuspend the beads by vortexing.
10. Briefly spin the tubes and leave them in a rack in the hood.

**9. Elution of the Final Library**

- Use 1.5 mL tubes, thermomix set to **25°C**, thermoblock set to **95°C**, and magnetic rack for 1.5 mL tubes.

1. Spin down with a microcentrifuge.
2. Pellet the beads using the magnetic rack and remove the supernatant.
3. Add **50 µL of TET buffer**.
4. Vortex and transfer the bead suspension to PCR 8-strip tubes.
5. Spin the strip briefly.
6. Incubate for **1 minute at 95°C** using the thermal cycler, followed by cooling to **25°C**.
7. Place the tubes into a magnetic rack.
8. Transfer the supernatant (final library) to fresh 1.5 mL low-bind tubes.
9. Freeze the library at -20°C in a DNA freezer.
10. **Break:** Take a break before proceeding to the next steps.

#### Protocol for Day 3 - qPCR and Indexing

**1. Prepare qPCR Standards**

**Dilute Libraries**

1. Prepare a **1:50 dilution** of each library in PCR 8-strip tubes:
    - **1 µL of library** + **49 µL of TET buffer**.
2. Vortex the mixtures and briefly spin them.

**Prepare Mastermixes**

3. Prepare two master mixes in 2 mL tubes:

**Mastermix A** (Measures total yield)

| Reagent | Volume (µL) | Total (for 50 reactions) (µL) |
|---|---|---|
| Mastermix (2x) | 12.5 | 625 |
| DEPC Water | 10 | 500 |
| IS7 (10 µM) | 0.5 | 25 |
| IS8 (10 µM) | 0.5 | 25 |

| Reagent | Volume (µL) | Total (for 50 reactions) (µL) |
|---|---|---|
| IS10 (10 µM) | 0.5 | 25 |
| **Total** | **24** | **1200** |

**Mastermix B** (Measures number of library molecules from spike-in control)

| Reagent | Volume (µL) | Total (for 5 reactions) (µL) |
|---|---|---|
| Mastermix (2x) | 12.5 | 62.5 |
| DEPC Water | 10 | 50 |
| IS7 (10 µM) | 0.5 | 2.5 |
| CL107 (10 µM) | 0.5 | 2.5 |
| CL118 (10 µM) | 0.5 | 2.5 |
| **Total** | **24** | **120** |

**Setup qPCR Plate**

4. Add **24 µL of Mastermix A** or **B** for each measurement to a well of an optical 96-well PCR plate. Both assays can be performed on one plate if 96 wells are not exceeded (or successively on two plates).
5. Add **1 µL of diluted library**, qPCR standard, or water to each well. **Note:** Do not discard the library dilutions; store at -20°C as a backup.
6. Seal the plate using optical caps or adhesive foil.
7. Vortex and briefly spin the plate in a plate centrifuge.

**2. Run PCR**

**PCR Program**

| Temperature (°C) | Time | Cycles |
|---|---|---|
| 95 | 10 min | |
| 95 | 30 sec | 45 |
| 60 | 30 sec | |
| 72 | 30 sec | |

8. Run the PCR with the program detailed above.
9. Determine the number of library and control molecules in each sample using the software provided with the qPCR system.

**3. Determine the Number of Cycles Needed for Indexing PCR**

**Calculation:**

| Reaction Volume (µL) | Input of DNA (µL) | Dilution |
|---|---|---|
| qPCR | 25 | 1:50 |
| Indexing PCR | 100 | 49 |
| | 4 | 49 |
| | 50 | |

| Optimal Cycle Numbers | Value |
|---|---|
| 4x the reaction volumes | 2.0 |
| 49x more sample DNA | measured cycles - 9 |
| 50x less diluted | ~11 cycles less |

**4. Indexing PCR**

**Prepare PCR Mastermix**

1. Prepare the PCR master mix in a 2 mL tube.

| Reagent | Volume (µL) | Total (for 17 reactions) (µL) |
|---|---|---|
| DEPC Water | 20 | 340 |
| AccuPrime Pfx Reaction Mix (x10) | 10 | 170 |
| AccuPrime Pfx Polymerase (2.5 U/µL) | 1 | 17 |
| **Total** | **31** | **527** |

2. After mixing, add primers and DNA:

| Reagent | Volume (µL) |
|---|---|
| P7_X Indexing Primer (10 µM) | 10 |
| P5_X Indexing Primer (10 µM) | 10 |
| Library (DNA) | 49 |
| **Total** | **100** |

**Add Primers**

3. Add the primer separately to each well so that each sample receives a unique pair of indices.

**5. Cycler Program**

| **Temperature (°C)** | **Time** | **Cycles** |
|---|---|---|
| Initial Activation | 95 | 2 min |
| Denaturation | 95 | 20 sec |
| Annealing | 60 | 30 sec |
| Elongation | 68 | 1 min |
| Final Elongation | 68 | 5 min |

**6. Purify PCRs**

1. Purify PCR products with the **MinElute PCR Kit** (see MinElute Protocol).
2. Elute the library in **20 µL TET buffer**.
3. Run samples on a **Bioanalyzer** for quality assessment.

#### Pooling

**Reagents Needed**

- HighPrep™ PCR solution
- 80% Ethanol
- Elution Buffer (TE Buffer)

**Important Preparation Step**

- **Bring the HighPrep™ PCR solution to room temperature for at least 30 minutes before use.**

**Procedure**

1. **Resuspend Beads:**
   - Shake the HighPrep™ PCR reagent thoroughly to fully resuspend the magnetic beads.
2. **Transfer Samples:**
   - Transfer the pooled samples into a **1.5 mL tube**.
3. **Add HighPrep™ PCR Reagent:**
   - Add **1.6x** the volume of HighPrep™ PCR reagent to the pooled samples.
4. **Mix Samples:**
   - Mix the HighPrep™ PCR reagent and the PCR sample by pipetting up and down **6-8 times**.
5. **Incubate:**

- Incubate the mixture for **5 minutes at room temperature**.

6. **Magnetic Separation:**
    - Place the tube on a magnetic rack for **3 minutes** or until the solution clears. The beads will be pulled to the side of the well.
7. **Remove Supernatant:**
    - While the tube is still on the magnet, remove and discard the supernatant by pipetting. **Caution:** Do not disturb the attracted beads while aspirating the supernatant!
8. **Ethanol Wash:**
    - With the tube on the magnet, add **200 µL of 80% ethanol** to each tube and incubate for **30 seconds at room temperature**.
9. **Remove Ethanol Supernatant:**
    - While the tube is still on the magnet, remove and discard the supernatant by pipetting.
10. **Repeat Ethanol Wash:**
    - Repeat steps 8-9 for a total of **two 80% ethanol washes**.
11. **Dry Beads:**
    - Dry the beads by incubating the tube for **10-15 minutes at room temperature** while still on the magnetic separation rack.

      **Note:** It is critical to completely remove all traces of alcohol but avoid over-drying the beads, as this will reduce yield.
12. **Remove from Magnetic Rack:**
    - Remove the tube from the magnetic separation rack.
13. **Add Elution Buffer:**
    - Add **40 µL of elution buffer** to each tube and pipette up and down **5 times** to mix. **Tip:** Prewarming the elution buffer at **55°C** can increase the yield.
14. **Incubate Again:**
    - Incubate for **2 minutes at room temperature**.
15. **Final Magnetic Separation:**
    - Place the tube back on the magnetic separation rack and wait **3 minutes** or until the magnetic beads clear from the solution.
16. **Transfer Eluate:**
    - Transfer the eluate (cleared supernatant) to a new tube for subsequent applications.
17. **Quality Check:**
    - Check the samples on a **BioAnalyzer** using the High Sensitivity Assay.

#### 1.2. Hybridization enrichment

##### 1.2.1. DNA baits

The DNA baits based enrichment was conducted according to IDT xGen™ hybridization capture of DNA libraries protocol with the AMPure XP Bead DNA concentration protocol option. A detailed outline of performed steps is provided in the following.

**Day 1: Hybridization Reaction**

**Hybridization Program Setup**

1. **Thermocycler Program:**
    - Set the lid temperature to 100°C.
    - Step 1: 95°C for 30 seconds.
    - Step 2: 63°C for 16 hours in a hybridization oven once the temperature is reached.

**Reagent Preparation**

1. **Thaw reagents:** Thaw all contents of the xGen Hybridization and Wash v2 Reagents to room temperature (RT).
    - Inspect the 2X Hybridization Buffer for any crystallization of salts. If crystals are present, heat the tube at 65°C, shaking intermittently until completely dissolved.
2. **Thaw xGen Hyb Panels:** Mix thoroughly and centrifuge briefly.

**Hybridization Mix Setup**

1. **Prepare the library mix:**
    - Add 500 ng of library to each tube containing Blocker components. If multiplexing samples, use 500 ng of each library.
    - Add 7.5 µL of Human Cot DNA.
    - Add 1.8X volume of AMPure XP beads.
    - Vortex thoroughly to mix and incubate for 10 minutes at room temperature.
2. **Magnet cleanup:**
    - Incubate the plate or tube on the magnet for at least 2 minutes or until the supernatant is clear.
    - Remove and discard the supernatant, then add 80% ethanol to cover the surface of the beads.
    - Incubate for 30 seconds, remove the ethanol, and repeat another wash.
    - Allow the beads to air dry for approximately 2 minutes, ensuring they do not over-dry.

**Hybridization Master Mix Preparation**

1. **Prepare the Hybridization Master Mix:**

- Mix the following components for each reaction:
  - xGen 2X Hybridization Buffer: 9.5 µL
  - xGen Hybridization Buffer Enhancer: 3 µL
  - xGen Universal Blockers (based on your library adapter): 2 µL
  - xGen Hyb Panel: 4.5 µL
- Total volume per reaction: 19 µL.

2. **Final preparation:** Vortex to mix and ensure the beads are fully resuspended. Incubate for 5 minutes at room temperature.
3. **Magnet separation:** Place on a magnet for 5–10 minutes or until the supernatant is clear.
4. **Transfer the supernatant:** Transfer 17 µL of the supernatant to a low-bind 0.2 mL PCR tube.
5. **Incubate and centrifuge:** Incubate for 5 minutes at room temperature, vortex, and briefly centrifuge.
6. **Start the HYB program** on the thermal cycler. Once the 63°C temperature is reached, place the samples in the incubation oven for hybridization.

**Day 2: Bead Capture and Wash**

**Bead Preparation**

1. **Prepare buffers:**
   - Ensure the Dynabeads M270 Streptavidin beads are at room temperature for at least 30 minutes before use.
   - Dilute the xGen buffers to create 1X working solutions as per the preparation table.
2. **Streptavidin bead wash:**
   - Vortex beads for 15 seconds, aliquot 50 µL of beads per capture, and add 100 µL of Bead Wash Buffer.
   - Pipette mix 10 times, place on the magnetic rack, and separate for 1 minute.
   - Discard the supernatant and repeat washing for a total of 3 washes.
   - Resuspend the beads in 17 µL per capture of Bead Resuspension Mix.

**Bead Capture Procedure**

1. **Transfer beads:** Add 17 µL of resuspended beads to the tube containing the hybridized sample.
2. **Incubation:** Place the sample in the thermal cycler for 45 minutes at 63°C, vortexing every 10–12 minutes.
3. **Washes:** Proceed to the heated washes using Wash Buffer 1 and Stringent Wash Buffer at 65°C, followed by room temperature washes with Wash Buffers 1, 2, and 3.

**Day 2: Post-Capture PCR and Clean-up**

**Post-capture PCR Setup**

1. **Amplification Mix Preparation:**
   - xGen 2x HiFi PCR Mix: 25 µL
   - xGen Library Amplification Primer Mix: 1.25 µL
   - Nuclease-Free Water: 3.75 µL
   - Total volume per reaction: 30 µL.
2. **Thermocycler Program:**
   - Polymerase activation: 98°C for 45 seconds.
   - Denaturation: 98°C for 15 seconds.
   - Annealing: 60°C for 30 seconds (13 cycles).
   - Extension: 72°C for 30 seconds.
   - Final extension: 72°C for 1 minute.
   - Hold at 4°C.

**Post-capture PCR Cleanup**

1. **AMPure XP bead cleanup:**
   - Add 1.5X volume of AMPure XP beads to each amplified sample and incubate for 5–10 minutes.
   - Wash twice with 80% ethanol, air dry beads for 1–3 minutes, and elute in 22 µL of Buffer EB.
   - Transfer 20 µL of eluate to a fresh tube ensuring no beads are carried over.

**Optional Step:**

- Store the purified PCR fragments per your laboratory's established procedures.

**Repeat the entire capture process for a second run if needed.**

##### 1.2.2. RNA baits

The RNA baits based enrichment was conducted according to myBaits v.5.02 manufacturer's manual. A detailed outline of performed steps is provided in the following.

**Day 1: Hybridization**

**Hybridization Mix Setup**

1. **Thaw and vortex**: Once the Hyb reagents have thawed, vortex them to homogenize and then briefly centrifuge.

2. **Assemble the Hybridization Mix**: Use a microcentrifuge (MC) tube, briefly vortex, and briefly centrifuge.

**Volumes adjusted for pipetting error:**

- **Hyb N**: 9.25 µL / rxn (46.25 µL total)
- **Hyb D**: 3.5 µL / rxn (17.5 µL total)
- **Hyb S**: 0.5 µL / rxn (2.5 µL total)
- **Hyb R**: 1.25 µL / rxn (6.25 µL total)
- **Water**: 1.1 µL / rxn (5.5 µL total); **Round 2:** 4.4 µL /rxn (22 µL total)
- **Baits**: 4.4 µL / rxn (22 µL total); **Round 2:** 1.1 µL / rxn (5.5 µL total)
- **Total**: 20 µL / rxn (100 µL total)

3. **Incubate**: At 60°C for 10 minutes in a heat block. Vortex occasionally. Remove from the heat block and let sit for 5 minutes.
4. **Aliquot**: For each capture reaction, aliquot 18.5 µL of Hybridization Mix to a 0.2 mL well/tube (HYBs).

**Blockers Mix Setup**

1. **Assemble the Blockers Mix**:
    - **Block O**: 2.5 µL / rxn (12.5 µL total)
    - **Block C**: 2.5 µL / rxn (12.5 µL total)
    - **Block X**: 0.5 µL / rxn (2.5 µL total)
    - **Total**: 5.5 µL / rxn (27.5 µL total)
2. **Aliquot**: For each capture reaction, aliquot 5 µL of Blockers Mix to a 0.2 mL well/tube.
3. **Add Libraries**: Add 7 µL of individual or pooled libraries to each Blockers Mix aliquot and mix by pipetting (LIBs).

**Reaction Assembly**

1. **Program Thermocycler**:
    - **Lid Temperature**: Set 5 to 10°C above each step temperature.
    - **Step 1**: 95°C for 5 minutes.
    - **Step 2**: 63°C, indefinite (hybridization temp).
2. **Load Thermocycler**: Put the LIBs in the thermal cycler, close the lid, and start the thermal program.
3. **Add HYBs**: Pause the program at hybridization temperature, put HYBs in the cycler, and resume.
4. **Mix**: After 5 minutes, pipette 18 µL of each HYB to each LIB and mix.
5. **Incubate**: Spin down LIBs, place in hybridization oven, and incubate overnight (24 hours).

**Day 2: Bind and Wash (Cleanup)**

**Washbuffer X**

1. **Prepare**: According to myBaits manual and sample volume.
2. **Heat**: Washbuffer X to hybridization temperature for 30 minutes.

**Bead Preparation**

1. **Aliquot Beads**: Pipette 150 µL beads into a low-bind 1.7 mL tube.
2. **Pellet and Wash**: Use magnet, remove supernatant, add 1000 µL Binding Buffer, vortex, centrifuge, and repeat wash three times.
3. **Final Resuspension**: Resuspend washed beads in 350 µL Binding buffer, aliquot 70 µL per reaction.

**Binding Beads and Hybrids**

1. **Heat Bead Aliquots**: Heat to 63°C for at least 2 minutes.
2. **Mix**: Transfer capture reactions to bead aliquots, mix, and incubate on thermomixer for 5 minutes. Centrifuge briefly.

**Bead Washing**

1. **Wash**: Pellet beads with magnet, discard supernatant, add 180 µL warmed Wash Buffer X, mix, centrifuge, and repeat wash three times.
2. **Resuspend**: After last wash, resuspend beads in Buffer E.

**Library Resuspension and Amplification**

1. **Resuspend Beads**: In 32 µL Buffer E.
2. **Incubate**: At 95°C for 5 minutes.
3. **Pellet and Collect**: Pellet beads, take supernatant.

**First Round PCR (14 cycles)**:

- **Master Mix (per rxn)**:
  - $H_2O$: 3.45 µL
  - Buffer 10X: 2.5 µL
  - dNTPs (25 mM): 0.25 µL
  - BSA: 1.0 µL
  - $MgSO_4$ (50 mM): 1.0 µL
  - HiFi (5 U/µL): 0.2 µL
  - IS5_bridge_P5 10 µM: 2.0 µL
  - IS6_bridge_P7 10 µM: 2.0 µL
- **Cleanup**: Bead Cleanup Kit, elute in 10 µL.

**Second Round PCR (8 cycles)**:

- **Master Mix (per rxn)**:
  - $H2O$: 3.45 µL
  - Buffer 10X: 2.5 µL
  - dNTPs (25 mM): 0.25 µL
  - BSA: 1.0 µL
  - MgSO4 (50 mM): 1.0 µL
  - HiFi (5 U/µL): 0.2 µL
  - IS5_bridge_P5 10 µM: 2.0 µL
  - IS6_bridge_P7 10 µM: 2.0 µL
- **Cleanup**: Bead Cleanup Kit, elute in 16-17 µL.

**Buffer Preparation**

- **Washbuffer X**:
  - **Hyb S**: 400 µL
  - **NF water**: 39.6 mL
  - **Wash Buffer**: 10 mL

### 2. Supplementary Results

#### 2.1. General data assessment

Information about DNA and RNA enriched data is shown in Table S1 The data set presents detailed information regarding the number of reads following trimming, the number of mapped reads, and the mapping outcomes of these reads to the mitochondrion and chloroplast for all samples. The samples are identified by library ID (libid), where "D" indicates DNA baits and "R" indicates RNA baits.

Following trimming, DNA baits yielded a total of 34.9 million sequences, while RNA baits produced 51.8 million sequences. Of these, 34.8 million (DNA) and 35.3 million (RNA) reads were successfully mapped to reference organelles. Specifically, RNA baits resulted in 6.41 million reads mapped to the mitochondrion and 28.9 million mapped to the chloroplast. In comparison, DNA baits yielded 6.27 million and 28.6 million reads mapped to the mitochondrion and chloroplast, respectively.

These findings highlight the greater efficiency of RNA baits in generating a higher volume of sequences and successfully mapping them to organelle references compared to DNA baits.

**Tab. S1: Summary of read mapping statistics for all samples. The table presents the number of reads after trimming, the total number of mapped reads, the number of reads mapped to the mitochondrial and chloroplast genomes, the number of unmapped reads, and the ability to construct consensus sequences for mitochondrial and chloroplast DNA.**

| libid | number_reads_after_trimming | number_of_mapped reads | mapped_to_mito | mapped_to_plastid | not_mapped | able_to_build_mito_consensus | able_to_build_plastid_consensus |
|---|---|---|---|---|---|---|---|
| PA014L-D | 2481755 | 2445859 | 408979 | 2036880 | 35896 | yes | yes |
| PA015L-D | 906457 | 948648 | 200058 | 748590 | NA | yes | yes |
| PA016L-D | 7278 | 6567 | 876 | 5691 | 711 | yes | yes |
| PA017L-D | 17452 | 16132 | 2390 | 13742 | 1320 | yes | yes |
| PA018L-D | 5056 | 4018 | 0 | 4018 | 1038 | no | no |
| PA019L-D | 52644 | 48591 | 7231 | 41360 | 4053 | yes | yes |
| PA020L-D | 66879 | 58329 | 5506 | 52823 | 8550 | yes | yes |
| PA021L-D | 15141 | 12808 | 1282 | 11526 | 2333 | yes | yes |
| PA022L-D | 9672 | 8471 | 1050 | 7421 | 1201 | yes | yes |
| PA023L-D | 128817 | 125159 | 23032 | 102127 | 3658 | yes | yes |
| PA024L-D | 2060906 | 2051213 | 361612 | 1689601 | 9693 | yes | yes |
| PA025L-D | 5231842 | 5303013 | 988245 | 4314768 | NA | yes | yes |
| PA026L-D | 60348 | 56131 | 8267 | 47864 | 4217 | yes | yes |
| PA027L-D | 2206545 | 2100753 | 379998 | 1720755 | 105792 | yes | yes |
| PA028L-D | 204884 | 191191 | 34900 | 156291 | 13693 | no | yes |
| PA029L-D | 2567 | 1972 | 0 | 1972 | 595 | yes | yes |
| PA030L-D | 247404 | 227028 | 38228 | 188800 | 20376 | yes | yes |
| PA031L-D | 158373 | 143570 | 24923 | 118647 | 14803 | yes | yes |
| PA032L-D | 916 | 702 | 0 | 702 | 214 | yes | yes |
| PA033L-D | 63704 | 56109 | 8987 | 47122 | 7595 | yes | yes |
| PA034L-D | 284842 | 261399 | 44142 | 217257 | 23443 | yes | yes |
| PA035L-D | 11449796 | 11808643 | 2252976 | 9555667 | NA | yes | yes |
| PA036L-D | 2312416 | 2332428 | 437668 | 1894760 | NA | yes | yes |
| PA037L-D | 2824696 | 2766467 | 471161 | 2295306 | 58229 | yes | yes |
| PA038L-D | 4102007 | 3850883 | 564556 | 3286327 | 251124 | yes | yes |
| PA039L-D | 19827 | 16992 | 1382 | 15610 | 2835 | yes | yes |
| **total_DNA_baits** | **3,49E+07** | **3,48E+07** | **6,27E+06** | **2,86E+07** | **5,71E+05** | | |
| PA014L-R | 5693413 | 4244580 | 934072 | 3310508 | 1448833 | yes | yes |
| PA015L-R | 2884561 | 2102210 | 508468 | 1593742 | 782351 | yes | yes |
| PA016L-R | 75663 | 45940 | 8092 | 37848 | 29723 | yes | yes |
| PA017L-R | 105217 | 65687 | 8686 | 57001 | 39530 | yes | yes |
| PA018L-R | 295784 | 185750 | 30142 | 155608 | 110034 | yes | yes |
| PA019L-R | 1843352 | 1196684 | 161462 | 1035222 | 646668 | yes | yes |
| PA020L-R | 3758206 | 2441009 | 422226 | 2018783 | 1317197 | yes | yes |
| PA021L-R | 649520 | 421753 | 55422 | 366331 | 227767 | yes | yes |
| PA022L-R | 119053 | 68912 | 12453 | 56459 | 50141 | yes | yes |
| PA023L-R | 879365 | 530091 | 98103 | 431988 | 349274 | yes | yes |
| PA024L-R | 5543956 | 3891352 | 721873 | 3169479 | 1652604 | yes | yes |

| | | | | | | | |
|---|---|---|---|---|---|---|---|
| PA025L-R | 9711975 | 7058290 | 1359989 | 5698301 | 2653685 | yes | yes |
| PA026L-R | 3756555 | 2427823 | 394962 | 2032861 | 1328732 | yes | yes |
| PA027L-R | 1217742 | 720979 | 118865 | 602114 | 496763 | yes | yes |
| PA028L-R | 183088 | 110993 | 17916 | 93077 | 72095 | yes | yes |
| PA029L-R | 3079 | 1770 | 328 | 1442 | 1309 | yes | yes |
| PA030L-R | 195083 | 115185 | 16751 | 98434 | 79898 | yes | yes |
| PA031L-R | 873515 | 518719 | 77637 | 441082 | 354796 | yes | yes |
| PA032L-R | 4390 | 2036 | 108 | 1928 | 2354 | yes | yes |
| PA033L-R | 288192 | 174948 | 27882 | 147066 | 113244 | yes | yes |
| PA034L-R | 1151696 | 716764 | 106502 | 610262 | 434932 | yes | yes |
| PA035L-R | 3495769 | 2317332 | 402991 | 1914341 | 1178437 | yes | yes |
| PA036L-R | 708727 | 528719 | 134923 | 393796 | 180008 | yes | yes |
| PA037L-R | 1694102 | 956882 | 178178 | 778704 | 737220 | yes | yes |
| PA038L-R | 6413220 | 4439320 | 600398 | 3838922 | 1973900 | yes | yes |
| PA039L-R | 82588 | 51891 | 7617 | 44274 | 30697 | yes | yes |
| **total_RNA_baits** | **5,18E+07** | **3,53E+07** | **6,41E+06** | **2,89E+07** | **1,65E+07** | | |

Table S2 presents the coverage data for all samples. Each sample (libid) includes data for both the mitochondrion and the chloroplast organelle. In general, the chloroplast show a higher total and average coverage compared to the mitochondrion across all samples.

**Tab. S2: Mapping coverage of for both organelles. Listed are the library id (libid) of the RNA-enriched dataset (R), the organelle type with corresponding total and average coverage.**

| libid | organelle | total_coverage | avg_coverage |
|---|---|---|---|
| PA014L-R | mitochondrion | 8601485 | 360513 |
| PA014L-R | chloroplast | 106478679 | 105445 |
| PA015L-R | mitochondrion | 6595599 | 256548 |
| PA015L-R | chloroplast | 42329947 | 390721 |
| PA016L-R | mitochondrion | 227368 | 119604 |
| PA016L-R | chloroplast | 1617218 | 102149 |
| PA017L-R | mitochondrion | 259666 | 905391 |
| PA017L-R | chloroplast | 2716112 | 984206 |
| PA018L-R | mitochondrion | 655785 | 614606 |
| PA018L-R | chloroplast | 5201966 | 762193 |
| PA019L-R | mitochondrion | 2868721 | 115674 |
| PA019L-R | chloroplast | 30084923 | 129906 |
| PA020L-R | mitochondrion | 4313964 | 594046 |
| PA020L-R | chloroplast | 37860820 | 116728 |
| PA021L-R | mitochondrion | 1458356 | 793879 |
| PA021L-R | chloroplast | 14615845 | 838786 |
| PA022L-R | mitochondrion | 344631 | 105715 |
| PA022L-R | chloroplast | 2326684 | 118088 |
| PA023L-R | mitochondrion | 2148419 | 176375 |
| PA023L-R | chloroplast | 15242275 | 266008 |

| | | | |
|---|---|---|---|
| PA024L-R | mitochondrion | 11466741 | 296291 |
| PA024L-R | chloroplast | 117833731 | 946836 |
| PA025L-R | mitochondrion | 18883428 | 441997 |
| PA025L-R | chloroplast | 192269853 | 152033 |
| PA026L-R | mitochondrion | 4853929 | 689087 |
| PA026L-R | chloroplast | 36561540 | 112837 |
| PA027L-R | mitochondrion | 2852131 | 111407 |
| PA027L-R | chloroplast | 24535117 | 242507 |
| PA028L-R | mitochondrion | 556295 | 49056 |
| PA028L-R | chloroplast | 4328562 | 733244 |
| PA029L-R | mitochondrion | 10643 | 107614 |
| PA029L-R | chloroplast | 60714 | 894168 |
| PA030L-R | mitochondrion | 515579 | 401447 |
| PA030L-R | chloroplast | 4513624 | 587673 |
| PA031L-R | mitochondrion | 1946727 | 333173 |
| PA031L-R | chloroplast | 15366228 | 292011 |
| PA032L-R | mitochondrion | 3230 | 123755 |
| PA032L-R | chloroplast | 76780 | 243129 |
| PA033L-R | mitochondrion | 808162 | 199842 |
| PA033L-R | chloroplast | 6364062 | 19033 |
| PA034L-R | mitochondrion | 2397129 | 262584 |
| PA034L-R | chloroplast | 21754874 | 395249 |
| PA035L-R | mitochondrion | 7657870 | 203299 |
| PA035L-R | chloroplast | 87004682 | 700651 |
| PA036L-R | mitochondrion | 3465111 | 402172 |
| PA036L-R | chloroplast | 18298358 | 232709 |
| PA037L-R | mitochondrion | 3763997 | 259532 |
| PA037L-R | chloroplast | 27686415 | 330304 |
| PA038L-R | mitochondrion | 6697646 | 437098 |
| PA038L-R | chloroplast | 95373882 | 145256 |
| PA039L-R | mitochondrion | 228294 | 173608 |
| PA039L-R | chloroplast | 1932286 | 148683 |

Table S3 shows the metadata on both sediment core samples, collected from the two locations, the EGB and the GOF as well as the reference data. Detailed are the core locations, depths, estimated ages, total organic carbon (TOC [%]) and associated climatic periods during the Holocene epoch. Key climatic events, such as the Medieval Climate Anomaly (MCA), Little Ice Age (LIA), and the Holocene Thermal Maximum (HTM), are highlighted alongside the prevailing estimated climatic conditions (e.g., cold or warm). Reference samples are also listed, denoted as mitochondrial and chloroplast. This dataset provides information about the paleoenvironmental and climatic evolution of the Baltic Sea region throughout the Holocene.

**Tab. S3: Meta data for all samples. Listed are the samples (libid), core and corresponding location, sampling sediment depth (depths), estimated ages in years cal BP (age(bp)),total organic carbon [%] (TOC)and associated climatic periods and phases during the Holocene epoch.**

| lib_id | core | location | depth | age(bp) | TOC | phases | climate_events | Holocene | Event/Period | Estimated_climatic_conditions |
|---|---|---|---|---|---|---|---|---|---|---|
| PA022L-R | EMB262_6_30_GC | EGB | 47 | 215 | 3,3 | modern Batic Sea | LIA | Late Holocene | LIA | cold |
| PA023L-R | EMB262_6_30_GC | EGB | 87 | 930 | 5,8 | modern Batic Sea | MCA | Late Holocene | MCA | warm |
| PA024L-R | EMB262_6_30_GC | EGB | 107 | 1063 | 7,4 | modern Batic Sea | MCA | Late Holocene | MCA | warm |
| PA025L-R | EMB262_6_30_GC | EGB | 112 | 1127 | 6,2 | modern Batic Sea | MCA | Late Holocene | MCA | warm |
| PA026L-R | EMB262_6_30_GC | EGB | 122 | 1257 | 7 | modern Batic Sea | MBS | Late Holocene | | warm |
| PA027L-R | EMB262_6_30_GC | EGB | 132 | 1395 | 5,5 | modern Batic Sea | LALIA | Late Holocene | LALIA | cold |
| PA028L-R | EMB262_6_30_GC | EGB | 162 | 1927 | 3,2 | modern Batic Sea | MBS | Late Holocene | | cold |
| PA029L-R | EMB262_6_30_GC | EGB | 182 | 2276 | 4,2 | modern Batic Sea | MBS | Late Holocene | | cold |
| PA030L-R | EMB262_6_30_GC | EGB | 262 | 3910 | 4,4 | modern Batic Sea | MBS | Late Holocene | | cold |
| PA031L-R | EMB262_6_30_GC | EGB | 282 | 4323 | 6,7 | Littorina Sea | LS | Mid-Holocene | | warm |
| PA032L-R | EMB262_6_30_GC | EGB | 292 | 4537 | 7,7 | Littorina Sea | LS | Mid-Holocene | | warm |
| PA033L-R | EMB262_6_30_GC | EGB | 302 | 4788 | 6,3 | Littorina Sea | LS | Mid-Holocene | | warm |
| PA034L-R | EMB262_6_30_GC | EGB | 322 | 5421 | 3,5 | Littorina Sea | HTM LS | Mid-Holocene | HTM | warm |
| PA035L-R | EMB262_6_30_GC | EGB | 342 | 5868 | 5,3 | Littorina Sea | HTM LS | Mid-Holocene | HTM | warm |
| PA036L-R | EMB262_6_30_GC | EGB | 422 | 7801 | 1,2 | Initial Littorina stage | HTM ILS | Mid-Holocene | HTM | warm |
| PA037L-R | EMB262_6_30_GC | EGB | 382 | 6801 | 4,9 | Littorina Sea | HTM LS | Mid-Holocene | HTM | warm |
| PA014L-R | EMB262_12_3_GC | GOF | 4 | -34 | 2,204 | modern Batic Sea | MWP | Late Holocene | MWP | warm |
| PA015L-R | EMB262_12_3_GC | GOF | 18 | 94 | 2,05 | modern Batic Sea | LIA | Late Holocene | LIA | cold |
| PA016L-R | EMB262_12_3_GC | GOF | 88 | 748 | 2,25 | modern Batic Sea | MCA | Late Holocene | MCA | warm |
| PA017L-R | EMB262_12_3_GC | GOF | 128 | 1122 | 2,375 | modern Batic Sea | MCA | Late Holocene | MCA | warm |
| PA018L-R | EMB262_12_3_GC | GOF | 148 | 1322 | 2,972 | modern Batic Sea | LALIA | Late Holocene | LALIA | cold |
| PA019L-R | EMB262_12_3_GC | GOF | 308 | 3482 | 2,08 | modern Batic Sea | MBS | Late Holocene | | cold |
| PA020L-R | EMB262_12_3_GC | GOF | 398 | 4272 | 2,008 | Littorina Sea | LS | Mid-Holocene | | cold |
| PA021L-R | EMB262_12_3_GC | GOF | 408 | 4359 | 2,178 | Littorina Sea | LS | Mid-Holocene | | cold |
| PA038L-R | EMB262_12_3_GC | GOF | 138 | 1213 | 2,492 | modern Batic Sea | MBS | Late Holocene | | warm |
| PA039L-R | EMB262_12_3_GC | GOF | 98 | 840 | 2,33 | modern Batic Sea | MCA | Late Holocene | MCA | warm |
| Sm_mitochondrion | Reference | SWO | | -60 | | modern Batic Sea | MWP | | | |
| Sm_plastid | Reference | SWO | | -60 | | modern Batic Sea | MWP | | | |

#### 2.2. RNA vs. DNA baits

DNA- and RNA baits achieved similar results, with RNA baits showing a higher resolution.

##### 2.2.1. Spatial differentiation

Our investigation of genetic diversity at the two sites, EGB and GOF, show that both DNA and RNA baits enriched samples depict comparable trends. However, the statistical significance of these trends varied based on the type of baits used for the enrichment. For the DNA baits enriched samples (Fig. Sn), we observed differences in genetic diversity between the two locations. Although these differences were not statistically significant (p-values: 0.052, 0.073, 0.116), they suggested a trend towards higher genetic diversity in EGB compared to GOF. In contrast, the RNA baits enriched samples (Fig. Sn) showed significant differences in genetic diversity between the two locations. EGB displayed a higher level of genetic diversity and higher diversity indices compared to GOF. These differences were statistically significant (p-values: 0.002, 0.005, 0.006), indicating distinct genetic structures in the populations of EGB and GOF.

In summary, our results highlight the importance of the type of samples used in genomic studies. Both DNA and RNA baits enriched samples reveal similar patterns of genetic diversity between the populations of EGB and GOF. Nevertheless, only the RNA baits enriched samples show statistically significant differences.

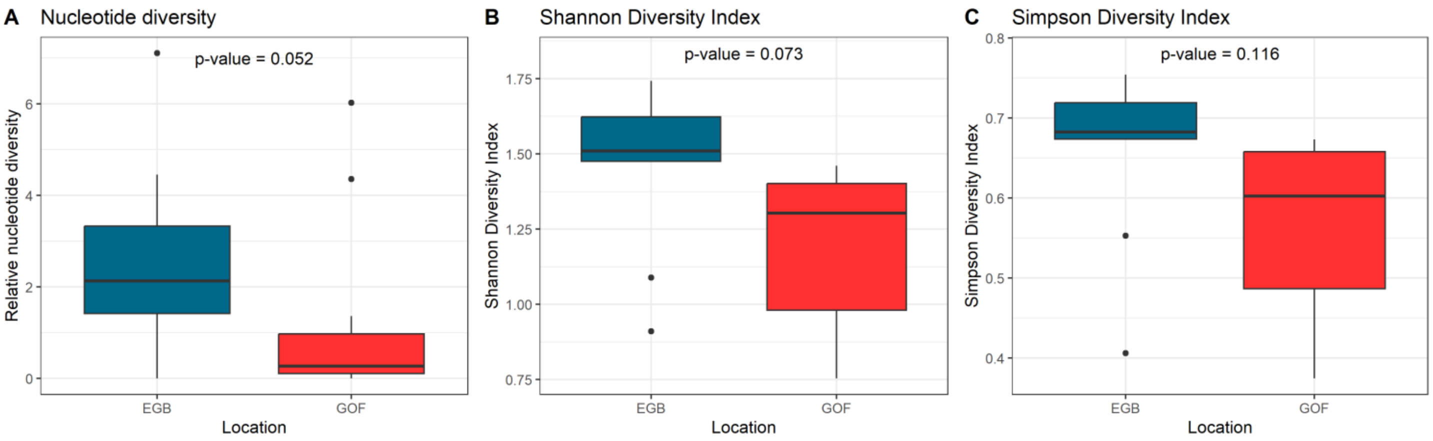


**Fig. S1: Comparative analysis of normalized nucleotide diversity and diversity indices of DNA baits enriched data. A) Nucleotide Diversity, B) Shannon, and C) Simpson) between EGB and GOF. Each bar graph shows data for both sites with p-values.**

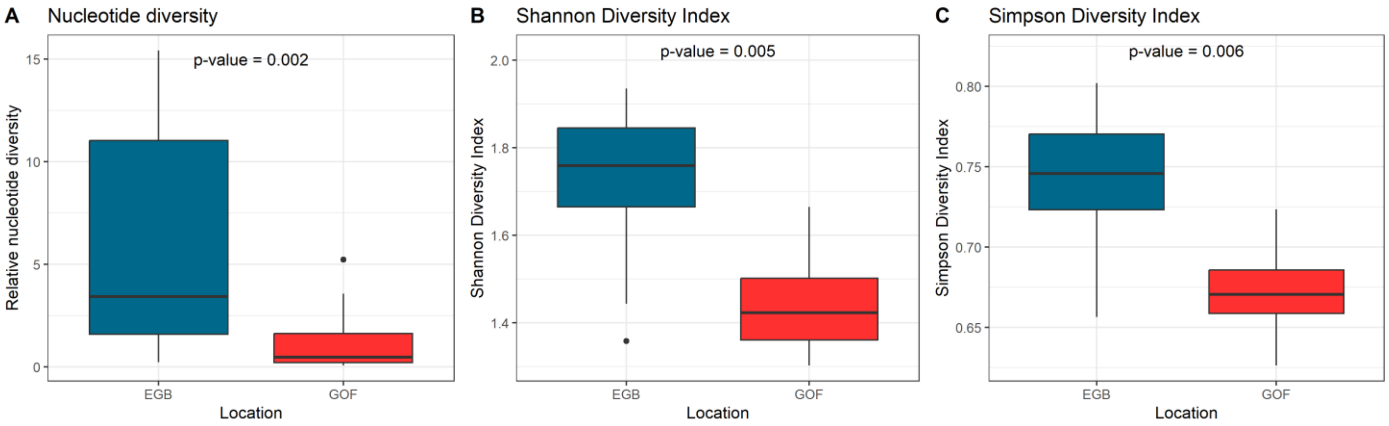


**Fig. S2: Comparative analysis of normalized nucleotide diversity and diversity indices of RNA baits enriched data. A) Nucleotide Diversity, B) Shannon, and C) Simpson) between EGB and GOF. Each bar graph shows data for both sites with significant p-values.**

##### 2.2.2. Climate events

The PCA of the allelic composition (Fig. 4) of *S. marinoi* organelles offers insights into how climate events might have influenced genetic variations at the two distinct locations: EGB and GOF. The PCA accounts for 39.1% of the variance in the DNA baits enriched data and 42.6% in the RNA baits enriched data.

The data from the DNA baits, presented in Figures 4A and 4C, demonstrate temporal changes. These changes appear to correspond with specific climate events, indicating that the genetic composition of the population is undergoing alterations in response to these occurrences. Figures 4B and 4D demonstrate analogous trends in the data from the RNA baits, thereby further that the population is adapting to environmental changes. The patterns observed using both DNA and RNA baits are comparable.

The clusters displayed in Figures 4C and 4D correspond to different climate events, including the Holocene Thermal Maximum (HTM), Little Ice Age/Late Antique Little Ice Age (LIA/LALIA), Medieval Climate Anomaly (MCA), and Medieval Warm Period (MWP). These clusters suggest a strong link between allelic variations and historical climate transitions. In the EGB, a longer time period is covered. We observe responses during the HTM, LALIA, MCA and LIA periods. However, the GOF location shows primarily changes during the LIA and MWP periods. It is important to note that the EGB samples do not cover the MWP. During periods of climatic stability, we observe more similar allelic compositions, which the population reverts to once an environmental change has settled or passed. This indicates a resilience over time and across climate events.

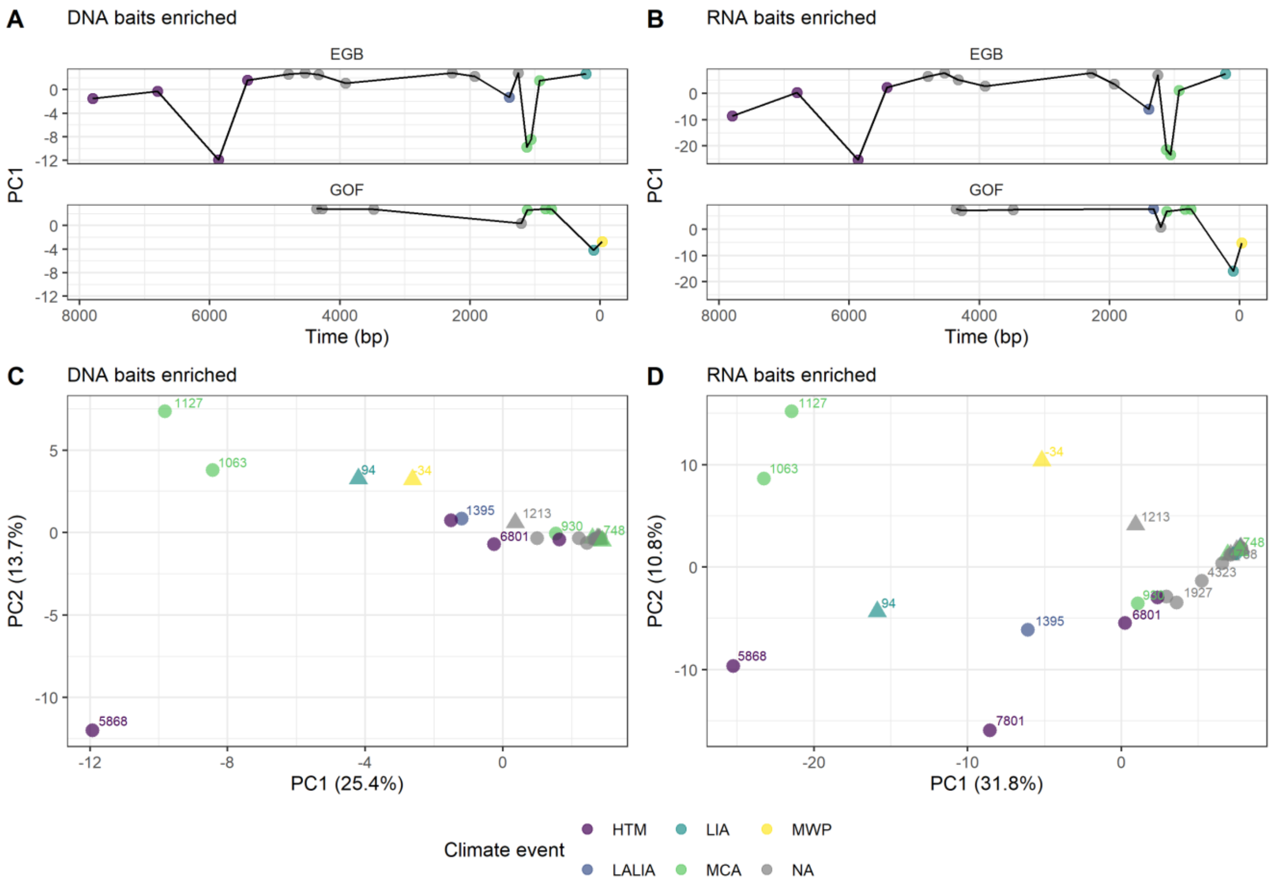


**Fig. S3: PCAs of allelic composition of *S. marinoi* organelles. PC1 against time (BP) in both locations A) DNA baits enriched data, B) RNA baits enriched data and PC1 & PC2 grouped by climate event and labeled by time (BP) C) DNA baits enriched data D) RNA baits enriched data. Climate events: HTM: Holocene Thermal Maximum, LALIA: Late Antique Little Ice Age, MCA: Medieval Climate Anomaly, LIA: Little Ice Age, MWP: Modern Warm Period.**

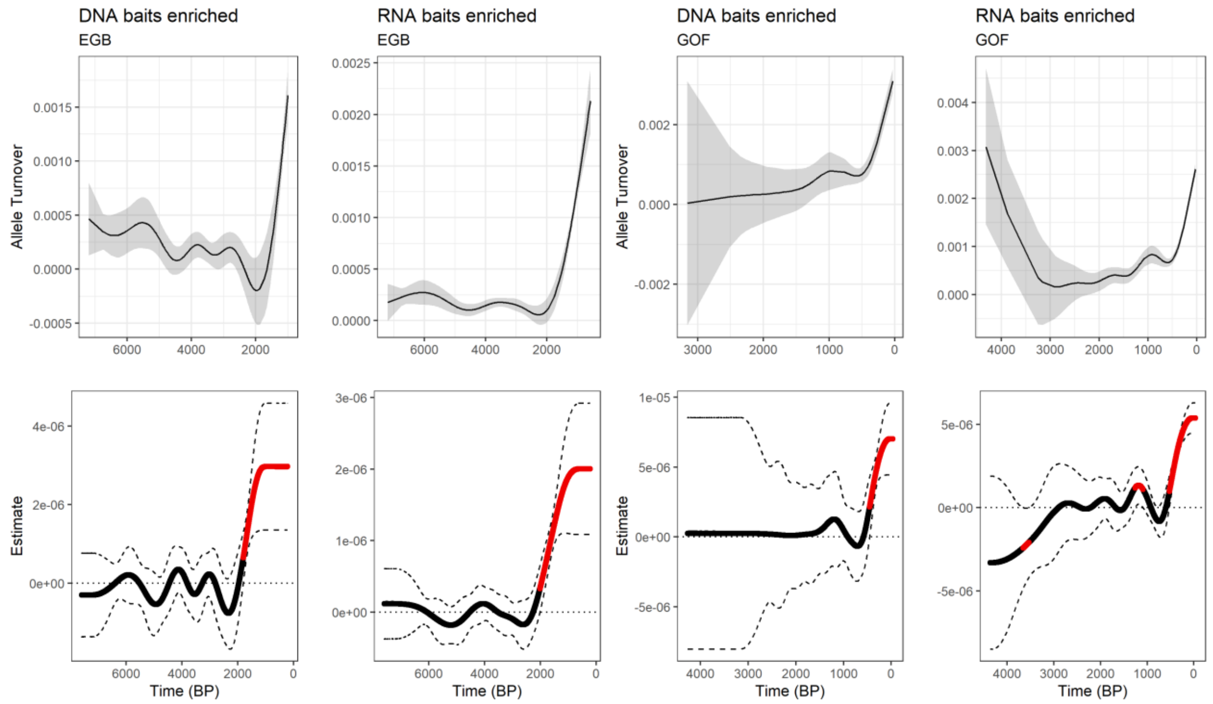


**Fig. S4: Analysis of allele turnover over time in *S. marinoi* population. The top row represents GAM models: The allele turnover variable was used to fit the GAM as a function of "age" (Time BP). The bottom row identifies periods of rapid change in allele turnover, determined by fitting a GAM and identifying periods where the first derivative of the fitted GAM significantly differs from zero. Data for both DNA baits and RNA baits enriched at both locations EGB and GOF are presented.**

#### 2.3. Influence of Damage pattern

In this study, we employed a principal component analysis (PCA) to investigate the relationship between allelic composition and climate events. To ascertain the influence of C-to-T substitution rates on composition, we conducted PERMANOVA and Pearson correlation tests, which yielded a statistically significant negative correlation between PC2 and the mean C-to-T substitution rate ($R = -0.508$, $p = 0.008$). However, no significant correlation was observed with PC1 ($R = -0.046$, $p = 0.823$). Furthermore, the PERMANOVA analysis indicated that the mean C-to-T substitution rate accounted for approximately 6.7% of the variation in the PCA space ($R^2 = 0.067$, $p = 0.19$), although this was not statistically significant. The PCA plot (Fig. S5) shows the distribution of samples based on their principal components (PC1 and PC2), with points colored according to the mean C-to-T substitution rate and shaped by location. Each point is labeled with the sample's age (in years BP), providing a temporal context.

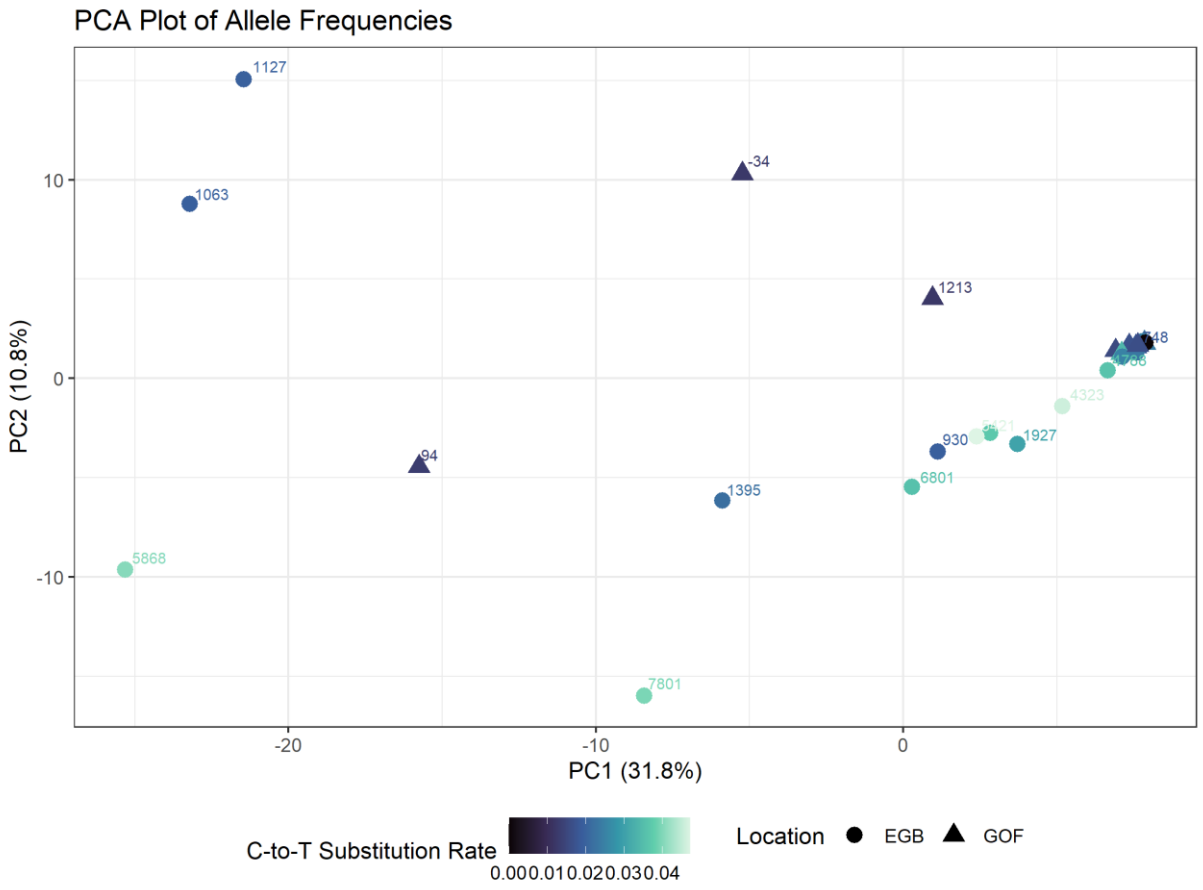


**Fig. S5: Principal component analyses of the allelic composition of *Skeletonema marinoi* organelles, demonstrating the genetic diversity and population structure. A) PC1 and PC2 are categorized by C-to-T-substitution rate and the age of each sample is given for both sites.**

#### 2.4. FST Analyses

The analysis of heterozygosity and FST over time between the two locations, EGB and GOF, provides insights into the regional differences. It should be noted that heterozygosity is not an appropriate measure of organelle diversity due to the influence of maternal inheritance. The FST values, which span approximately from 0 to 0.1, indicate a lack of discernible genetic differentiation between the two locations. This implies that the populations from EGB and GOF are not markedly disparate in terms of genetic structure (Fig. Sx).

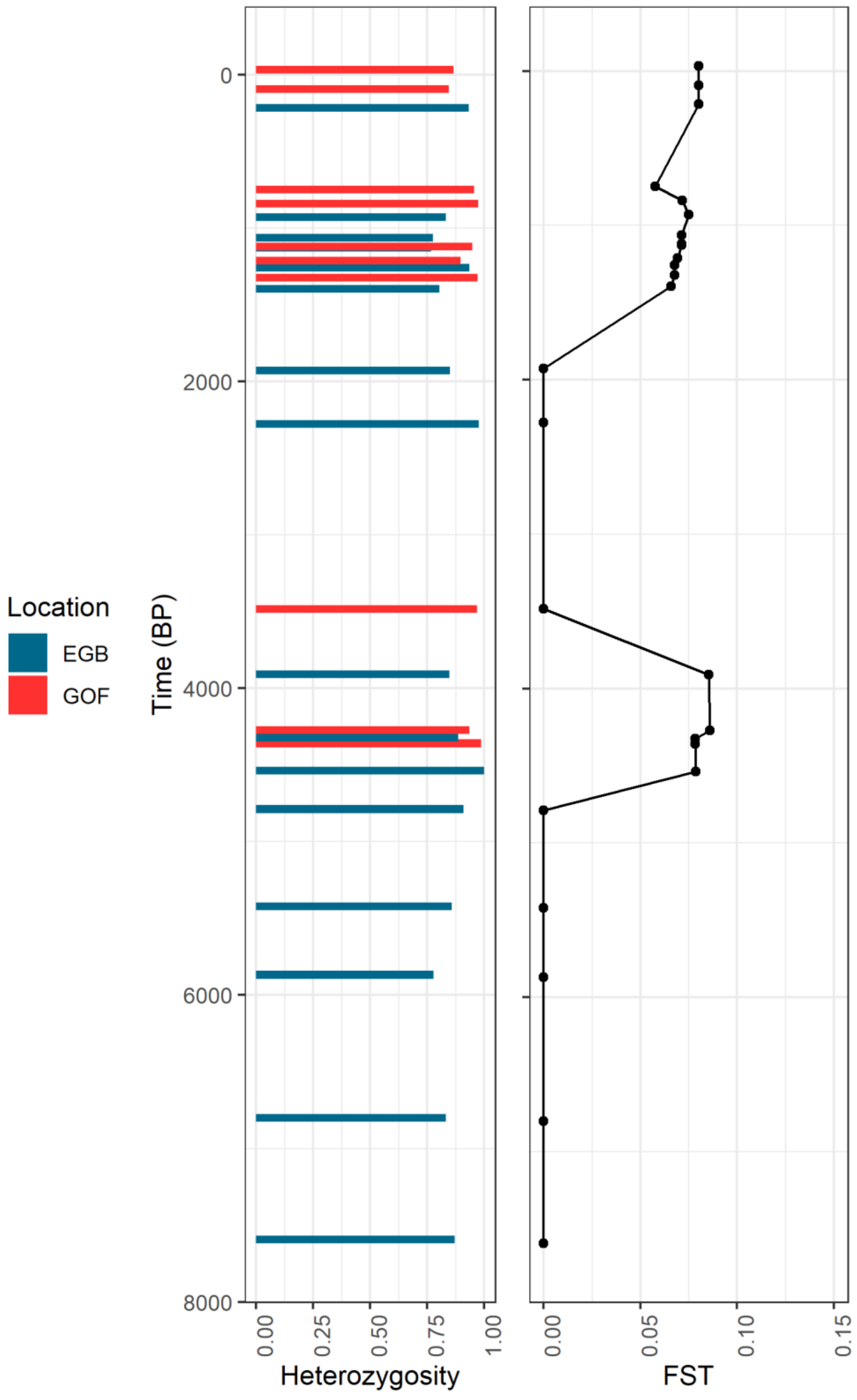


**Fig. S6: Comparative analysis of heterozygosity (left) and FST (right) over time between two locations, EGB and GOF.**

#### 2.5. Nucleotide Diversity

The analysis of nucleotide diversity (pi) across different time periods revealed significant differences when comparing the beginning of the modern Baltic Sea period with both the Littorina Sea and Roman-Hanseatic periods. Specifically, the following observations were made:

The mitochondrion data revealed a significant higher relative pi during the beginning Modern Baltic Sea period (2000-4000 cal BP) compared to the Littorina Sea period (more than 4000 cal BP) ($p = 0.026$). Significant differences were observed between the beginning of the Modern Baltic Sea period (4000-2000 cal BP) and the Littorina Sea period (until 4000 cal BP) for chloroplast data ($p = 0.017$). Furthermore, significant differences were observed between the Roman-Hanseatic period (2000-100 cal BP) and the beginning of Modern Baltic Sea period (4000-2000 cal BP) ($p = 0.029$). It is important to note that all periods after the Littorina Sea are considered part of the Modern Baltic Sea (4000 cal BP until Present).

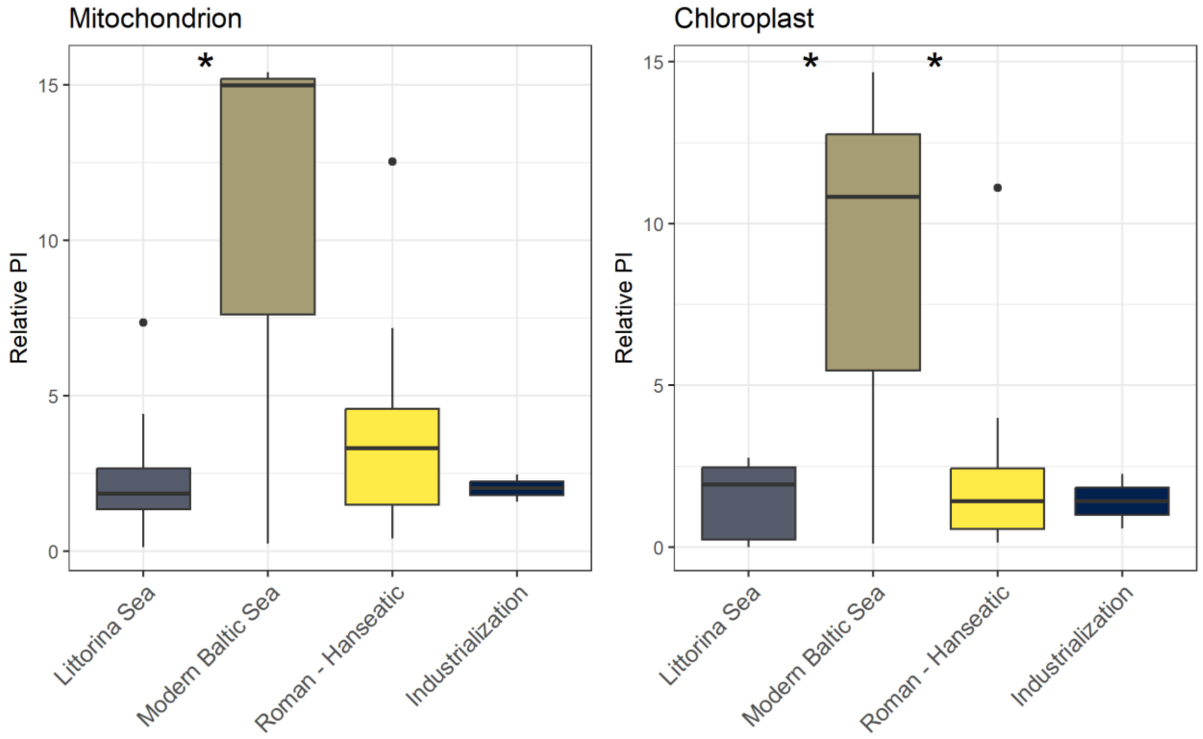


**Fig. S7: Box plots depicting relative nucleotide diversity (pi) across four historical periods for mitochondrion and chloroplast DNA sequences. Asterisks indicate statistically significant differences between groups based on Tukey's post-hoc test (*: $p < 0.05$). Littorina Sea (until 4000 BP), Modern Baltic Sea (4000-2000 BP), Roman-Hanseatic (2000-100 BP), Industrialization (100 BP until Present).**
